# Proximal termination generates a transcriptional state that determines the rate of establishment of Polycomb silencing

**DOI:** 10.1101/2023.07.06.547969

**Authors:** Govind Menon, Eduardo Mateo-Bonmati, Svenja Reeck, Robert Maple, Zhe Wu, Robert Ietswaart, Caroline Dean, Martin Howard

## Abstract

Chromatin-mediated transcriptional silencing by Polycomb Repressive Complex 2 (PRC2) is critical for gene regulation in development and environmental responses. However, the mechanism and timescales controlling de novo establishment of PRC2 silencing are unclear. Here, we investigate PRC2 silencing at Arabidopsis *FLOWERING LOCUS C* (*FLC*), known to involve co-transcriptional RNA processing, histone demethylation activity, and PRC2 function; but so far not mechanistically connected. We develop and then test a computational model that describes how proximal polyadenylation/termination mediated by the RNA binding protein FCA induces H3K4me1 removal by the histone demethylase FLD. H3K4me1 removal feeds back to reduce RNA Pol II processivity and thus enhance early termination, thereby repressing productive transcription. The model predicts that this transcription-coupled repression controls the level of transcriptional antagonism to Polycomb action, Thus, the effectiveness of this repression dictates the timescale for establishment of Polycomb H3K27me3 silencing. Experimental validation of these model predictions allowed us to mechanistically connect co-transcriptional processing to setting the level of productive transcription at the locus, which then determines the rate of the ON to OFF switch to PRC2 silencing.

## Introduction

Transcriptional regulation is often mediated by chromatin states that are key to developmental changes and environmental responses in many organisms. These chromatin states are characterised by histone modifications, and trans-factors that recognise these modifications, which together influence the level of productive transcription. Such chromatin states can be heritable across DNA replication and mitosis, with controlled switching of key developmental loci between active and silenced states underpinning the changes in gene expression programs that drive differentiation.

Polycomb Repressive Complex 2 (PRC2) mediated H3K27me3 is a system that generates such heritable and switchable chromatin states. The mechanisms enabling inheritance in this case include read-write reinforcement of existing histone modifications and antagonism between transcriptional activity and silencing modifications (Berry et al., 2017; Dodd et al., 2007; Holoch et al., 2021). Such mechanisms can generate bistability producing all or nothing (ON or OFF) transcriptional states (Berry et al., 2017; Pease et al., 2021). Each gene copy can be stably maintained in either an ON or OFF state even in dividing cells and copies can be independently switched between states. Such switching has been achieved through experimental perturbations in various studies (Holoch et al., 2021), but there are few examples where the mechanisms that enable the set-up and switching of these states are studied in natural developmental contexts.

One such example is PRC2-mediated regulation at the Arabidopsis floral repressor gene *FLOWERING LOCUS C* (*FLC*) (Menon et al., 2021). The winter cold-induced epigenetic silencing is an ON/OFF cell-autonomous, stochastic switching of individual *FLC* copies into a Polycomb silenced state (Fig. 1A) (Angel et al., 2011; Berry et al., 2015; Csorba et al., 2014; Lövkvist and Howard, 2021; Menon et al., 2021). In this process, the silencing histone modification H3K27me3 is first nucleated (deposited at a specific intragenic site) during the cold, followed by spreading of H3K27me3 across the locus through a cell-cycle dependent mechanism during growth after cold (Yang et al., 2017). Establishment of Polycomb silencing requires specific Polycomb accessory proteins and chromatin remodellers, some specific to the nucleation or spreading phases.

**Figure 1:**
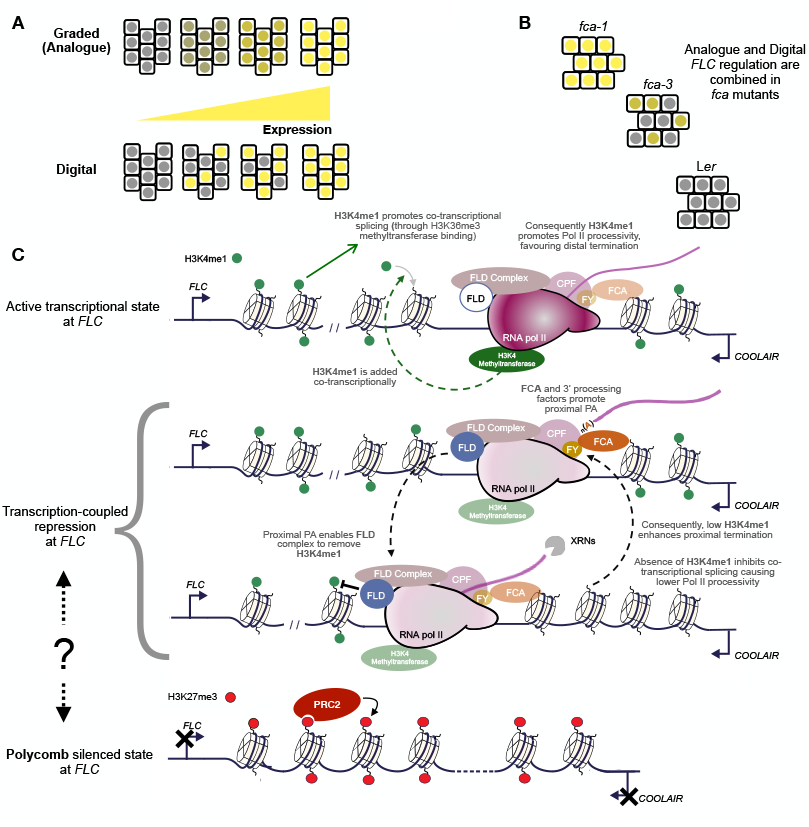
**(A)** Schematic showing two possible modes of regulating population level expression – Analogue and Digital. **(B)** Schematic of combined digital and analogue regulation observed experimentally in *fca* mutants. **(C)** Schematic showing the transcription coupled repression mechanism at *FLC*. Activating H3K4me1 can be added during productive transcription (green dashed arrow). The repression mechanism consists of proximal polyadenylation and termination of sense and antisense transcription mediated by the RNA binding protein FCA, which then enables removal of transcriptionally activating H3K4me1 across the locus by the putative histone demethylase FLD. The removal of H3K4me1 can inhibit co-transcriptional processing by preventing the addition of H3K36me3, and consequently promotes further proximal termination. Thus, a feedback loop (highlighted by the black dashed arrows) is formed, where the occurrence of proximal termination promotes further proximal termination. Also shown is a schematic of the Polycomb silenced state at *FLC*, but how this is linked to transcription-coupled repression has been unclear. Supplementary file: Movie_1 contains an animation summarising the processes and interactions shown here.

In rapid-cycling Arabidopsis accessions *FLC* silencing is constitutive, occurring early in development. This silencing requires the RNA binding protein FCA, which interacts with Cleavage and Polyadenylation Specificity Factor (CPF) machinery (Simpson et al., 2003). FCA promotes proximal termination of *FLC* sense transcription in the developing embryo (Schon et al., 2021), and of *FLC* antisense transcription in the seedling (Liu et al., 2010). *FLC* silencing also requires FLD (Liu et al., 2007), a plant homologue of the conserved mammalian histone K4 demethylase LSD1 (Martignago et al., 2019). The role of FCA and other co-transcriptional regulators in this mode of *FLC* silencing, and the observed FCA interactions with termination and 3’ processing factors, suggests a transcription-coupled mechanism that links to Polycomb silencing. Indeed, the accompanying paper identifies a role for the CPF phosphatase module, a key regulator of RNA Pol II termination, in this mode of *FLC* silencing (Mateo-Bonmatí et al., 2023). This mechanism thus has the potential to provide a window into the link between transcription-coupled repression and Polycomb silencing.

Transcription-coupled mechanisms have been implicated in the establishment of heterochromatin in many organisms, through 3’ processing and non-coding RNA. A recent study indicates a wide-ranging role for 3’ processing in heterochromatin establishment in *S. pombe* (Shimada et al., 2021). For example, Clr4-mediated heterochromatin assembly by the RNAi-independent pathway involves non-canonical polyadenylation and 3’ processing factors (Chalamcharla et al., 2015; Tucker et al., 2016; Vo et al., 2019). Transcription termination at non-canonical sites mediated by the conserved cleavage and polyadenylation factor (CPF) and the RNA-binding protein Mmi1 have been shown to be necessary for Clr4-driven assembly of facultative heterochromatin at meiotic and environmental response genes. Another RNA binding protein, Seb1, has been shown to promote heterochromatin formation, through a mechanism linked to termination of noncoding transcription (Marina et al., 2013; Wittmann et al., 2017). For PRC2-mediated silencing, early termination has been suggested to reduce nascent RNA inhibition of PRC2, by limiting the physical length of the transcript produced (Kaneko et al., 2014). Nevertheless, the exact mechanisms linking co-transcriptional processing to chromatin regulation are still poorly understood.

Exploiting an allelic series of *fca* mutations, we recently showed that the FCA-mediated mechanism can generate a graded regulation of *FLC* expression (Antoniou-Kourounioti et al., 2023). Thus, *FLC* can be regulated both in a graded manner (or *analogue* regulation) or by an ON/OFF mode of regulation (or *digital* regulation) as in the cold-induced silencing (Fig. 1A). The intermediate *fca-3* allele revealed that FCA-mediated analogue regulation precedes an ON/OFF developmental switch at *FLC* (Fig. 1B)(Antoniou-Kourounioti et al., 2023). Thus, the study highlighted the switch from analogue to digital regulation but did not mechanistically address HOW that switch occurred. In this study, we therefore asked: How can co-transcriptional processing mediated by FCA and the linked function of the chromatin modifier FLD achieve the observed *analogue* mode/graded manner of *FLC* regulation? Does the observed ON to OFF switch involve PRC2? If so, what is the mechanistic link between this analogue transcription-coupled mode of regulation and a Polycomb ON/OFF *digital* mode of regulation? Specifically, how is an FCA/FLD mediated *repressed* state (with enhanced early termination of transcription enabling graded regulation of the amount of full length transcript produced) linked to a PRC2 mediated *silenced* state (OFF state with a low frequency of transcription initiation)?

Using mathematical modelling combined with molecular approaches, we dissect the interactions between co-transcriptional processing and chromatin regulation that underpin how transcription-coupled repression can lead to Polycomb-mediated silencing of *FLC*. Specifically, the model describes sense and antisense transcription, alternative transcriptional termination, the dynamics of activating (H3K4me1) and silencing (H3K27me) modifications, and the interactions between these components (Fig. 1C). Importantly, the complexity of these feedbacks could not be interpreted without the use of modelling approaches. We first showed that the model can successfully generate intermediate transcriptional outputs at non-Polycomb silenced *FLC* copies (analogue/graded regulation), while also allowing Polycomb nucleation to occur slowly over time (ON/OFF switch). We then used the model to qualitatively predict changes in activating H3K4me1 and silencing H3K27me3 modifications at *FLC* over time, and experimentally validated these predictions by measuring changes in these modifications over a developmental time-course, showing further that the model can be quantitatively fit to capture these changes. Further validation was provided by measuring timecourse changes in *FLC* expression, where the model parameter fit obtained from histone modification changes also achieves a satisfactory quantitative match with no additional changes. The separate roles of the co-transcriptional repression and Polycomb silencing as predicted by the model are further validated by examining a mutant of a Polycomb methyltransferase. The model clearly elaborates how FCA/FLD-mediated transcription-coupled repression, induced by co-transcriptional processing steps as described in the accompanying paper (Mateo-Bonmatí, 2023), can influence the timing of the digital switch to Polycomb silencing. The combination of molecular genetics and the theoretical modelling described here provide a quantitative description of how co-transcriptional processing can quantitatively set transcriptional states that determine rates of Polycomb silencing, a mechanism that is likely to be generally relevant in many organisms.

## Results

### Molecular framework governing the *FLC* transcription-coupled chromatin repression pathway

The RNA binding protein FCA promotes proximal polyadenylation/termination at many targets in the Arabidopsis genome (Sonmez et al., 2011). FCA is known to directly interact *in vivo* with FY, a CPF component needed for silencing homologous to yeast Pfs2 and mammalian WDR33 (Simpson et al., 2003). FCA also indirectly interacts with other cleavage and polyadenylation specificity factors (Fang et al., 2019), as well as the 3’ processing factors CstF64 and CstF77 (Liu et al., 2010). FCA-mediated silencing also requires the involvement of CDKC2, a component of the transcription elongation factor P-TEFb (Wang et al., 2014), as well as the core spliceosome component PRP8 (Marquardt et al., 2014), suggesting it is the process of transcription, co-transcriptional processing, and proximal termination, rather than just the transcripts themselves, that drive silencing. FCA is proposed to recognize any situation where RNA Pol II has stalled and transcription termination is required to clear a ‘tangle’ in the chromatin (Baxter et al., 2021; Xu et al., 2021).

*FLC* silencing mediated by FCA requires the protein FLD, a homologue of the mammalian H3K4/K9 demethylase LSD1 (Liu et al., 2007). FLD is thought to demethylate H3K4me1 in Arabidopsis (Inagaki et al., 2021; Martignago et al., 2019). FLD has been shown to associate *in vivo* with the SET domain protein SDG26, which interacts with FY, indicating a physical link between processing of antisense transcripts and chromatin remodelling (Fang et al., 2020). FLD also physically associates in vivo with LD, a TFII-S domain containing protein and homolog of the CPF phosphatase subunit, PNUTS in human and Ref2 in yeast. This strong association of chromatin modifiers such as FLD with RNA Pol II regulators shown in the accompanying paper, identifies a mechanism where FCA-mediated proximal cleavage and polyadenylation triggers FLD-mediated removal of H3K4me1 (Mateo-Bonmatí, 2023).The Arabidopsis H3K36 methyltransferase SDG8 is known to recognise H3K4me1, so H3K4me1 removal might be expected to affect H3K36 methylation. *FLC* upregulation in an *fca* mutant is rescued in an *sdg8 fca* double mutant (Fang et al., 2020), indicating that H3K36me3 may be essential to set up an active transcriptional state at *FLC*. This is consistent with our previous ChIP-qPCR data at *FLC*, showing high levels of H3K4me1 and H3K36me3 at the locus in a high transcriptional state and low levels of both these modifications in a low transcriptional state (Fang et al., 2020). Recent evidence from *Saccharomyces cerevisiae*, shows that H3K36me3 is required for the recruitment of splicing factors and effective co-transcriptional splicing (Leung et al., 2019). Based on this evidence, FLD-mediated removal of H3K4me1 may cause reduced H3K36 methylation, thus inhibiting co-transcriptional splicing, consequently reducing Pol II processivity and promoting further early transcription termination. The FLD mechanism would thus form a feedback loop where early termination enhances itself through H3K4me1 removal. The above molecular mechanisms and possible feedback interaction are summarised in the schematic shown in Fig. 1C.

### FCA promotes premature termination of both sense and antisense transcription at *FLC*

The *FLC* sense proximal transcripts are detectable in young embryos (Schon et al., 2021) but less so in vegetative tissue, likely due to rapid nascent RNA turnover. In contrast, the *FLC* antisense RNAs transcribed from the 3’ end of *FLC* (collectively called *COOLAIR*), which are alternatively polyadenylated (PA) at a proximal PA site within the *FLC* gene or a distal PA site in the vicinity of the *FLC* promoter (Liu et al., 2010; Swiezewski et al., 2009), are both readily detectable. The mechanism underlying the differential turnover of proximally polyadenylated transcripts is currently unknown, but differential fates of coding and non-coding transcripts are not uncommon (Nojima and Proudfoot, 2022). To assess the relative roles of FCA-mediated premature termination of sense/antisense transcripts in silencing, we compared repression of a wild-type *FLC* transgene (*FLC-15*), with that of an *FLC* transgene where the antisense promoter has been replaced (*TEX 1*.*0)* altering the type and quantity of antisense transcript production (Csorba et al., 2014; Marquardt et al., 2014; Wang et al., 2014). We performed the comparison in two genetic backgrounds: one with the endogenous FCA carrying a null mutation (*fca-9* mutant in *Col-0*), thus having no functional FCA, and the other with the same mutant of the endogenous FCA, but carrying a transgenic 35S::FCA construct (Liu et al., 2007), giving FCA overexpression. We made two observations (Fig. 2A): (i) the wild-type *FLC* transgene is significantly repressed by FCA overexpression (as previously reported (Liu et al., 2007)); (ii) this FCA dependent repression of the wild-type *FLC* transgene is significantly attenuated ∼10 fold in *TEX 1*.*0*, but repression is not lost completely. Thus, the antisense transcription mechanism only partially accounts for *FLC* repression; a fact supported by a CRISPR deletion of the antisense promoter at the endogenous *FLC* locus causing a small *FLC* upregulation (Zhao et al., 2021). To further explore the importance of FCA-mediated proximal termination we analysed production of sense polyadenylated *FLC* transcripts by Quant-seq (Moll et al., 2014) in the *fca* allelic series (Antoniou-Kourounioti et al., 2023; Koornneef et al., 1991).Three genotypes give different levels of *FLC* expression at the whole plant level: the wild-type L*er*, with low *FLC* expression, the *fca-1* mutant, with high *FLC* expression, and the *fca-3* mutant, with intermediate *FLC* expression. Proximally polyadenylated transcripts for the *FLC* transcript were detected for all genotypes analysed at sites close to those reported in embryos (Fig. 2B) (Schon et al., 2021). Altogether these data support a model whereby FCA-mediated proximal termination of both sense and antisense transcription contributes to repression of *FLC* transcription.

**Figure 2:**
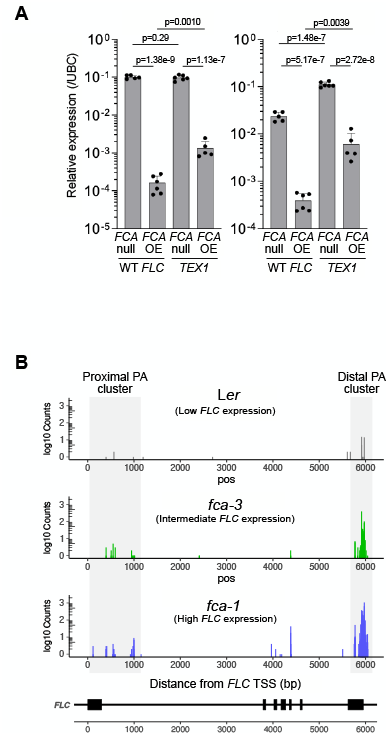
**(A)** qPCR data for *FLC* expression, showing repression of a transgenic wild-type *FLC* construct (*FLC-15* indicated by WT *FLC*) in an FCA overexpression genotype (*35S::FCA* indicated by *FCA* OE), in 14 day old seedlings, and the attenuation of this repression for an *FLC* transgenic construct where antisense transcription is disrupted (*TEX1*). All genotypes shown are in an *fca-9* (*FCA* null mutant) background. Error bars represent mean +/- s.e.m. p-values are shown for a statistical comparison of the mean values using the Student’s t-test. **(B)** Quantseq data for 7-day old seedlings (7 days after sowing), showing quantification of polyadenylated transcripts mapped to the *FLC* locus in the parental Landsberg (L*er*) genotype and the *fca* mutants. Individual replicates are shown in Fig. S4D.

### Computational model of *FLC* chromatin silencing through alternative termination

The silenced state of *FLC* is associated with H3K27me3 coverage of the whole locus (Wu et al., 2016). Genetic analysis also shows that the plant Polycomb methyltransferase CURLY LEAF (CLF) is essential for FCA-mediated silencing of *FLC* (Fang et al., 2020). However, despite the detailed molecular data on 3’ processing-mediated silencing described above, and in the accompanying paper (Mateo-Bonmatí, 2023), how the various elements linked to Polycomb silencing was unknown, due in part to the complex feedbacks involved. To bridge this gap, we therefore constructed a computational model based on the above features (Fig. 3A), using our prior knowledge of Polycomb dynamics known to generate digital ON/OFF memory states. We then used this model to dissect the underlying mechanism of the transition elucidating how digital silencing might be integrated into FCA-mediated 3’ processing. Important features of the new model are described below:

**Figure 3:**
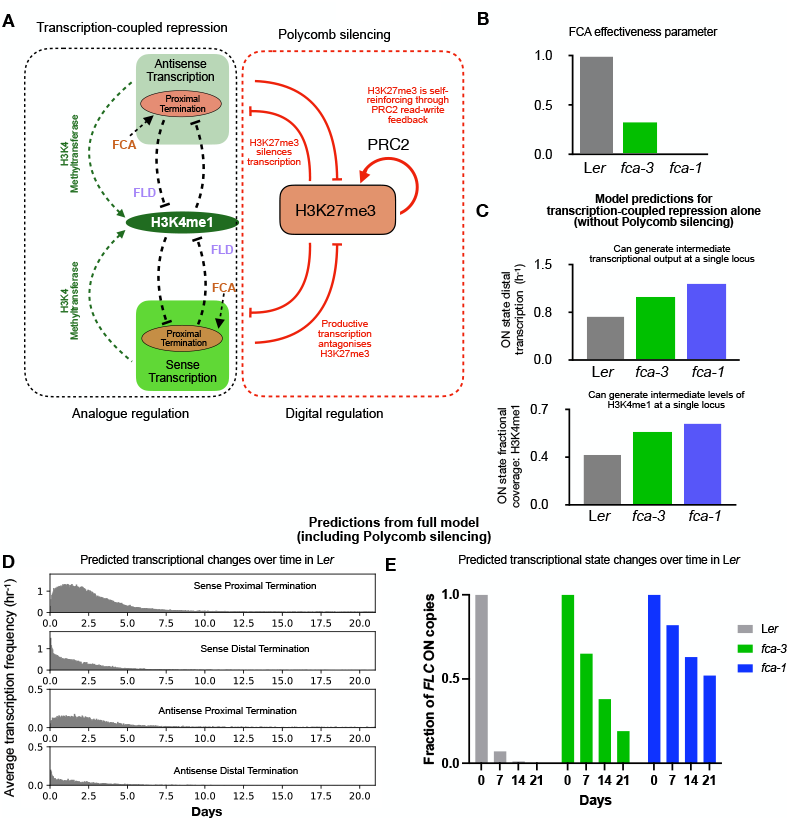
**(A)** Schematic of the combined mathematical model incorporating the transcription-coupled repression mechanism and PRC2 mediated silencing. The schematic highlights the key processes and interactions captured by the mathematical model. For the transcription coupled repression mechanism (as depicted in Fig. 1C) this includes sense and antisense transcription, co-transcriptional addition of H3K4me1 (green dashed arrows), FCA mediated proximal termination of transcription, FLD mediated removal of H3K4me1, and the promotion of proximal termination by low H3K4me1 coverage. Feedback is again highlighted by black dashed lines. For PRC2 silencing the model includes H3K27 methylation, PRC2 mediated read-write maintenance of these modifications, and mutual antagonism between H3K27me3 and transcription. **(B)** FCA effectiveness parameter values for the three genotypes. **(C)** Top panel: Model predicted frequency of distal transcription events (steady state average over 1000 simulated loci) in an ON state, for the model of the transcription-coupled repression mechanism shown in (A) but without the Polycomb silencing module. Each simulated locus was started in an active state with full H3K4me1 coverage and allowed to equilibrate over a duration of 5 cell cycles. Averaging was performed over the duration of the following 10 cell cycles. Bottom panel: Model predicted fractional coverage of H3K4me1 in an ON state from the above simulations. **(D)** Predicted transcriptional changes (quantified by frequency of transcription events) over time obtained by simulating the combined model shown in (A). Frequencies of the different transcription events are averaged over 1000 simulated copies of the *FLC* locus, all started in an active state with full H3K4me1 coverage. **(E)** Predicted changes in the fraction of ON copies in the *fca* mutants, computed from simulations of 1000 copies over 30 cell cycles, started in a full H3K4me1 covered ON state. Fractions are shown at the starting timepoint (0), and at the end of 7, 14, and 21 days from the start. See Supplementary Information for the definition of ON and OFF states used in this analysis.

The model includes both activating (H3K4me1) and repressive (H3K27me3) modifications at each histone across the *FLC* locus, assumed mutually exclusive on the same H3 tail, consistent with the reported mutual exclusivity of these modifications (Shema et al., 2016). The model also incorporates sense and antisense transcription at the locus. Importantly, the model assumes distinct roles for the two kinds of modifications in transcription regulation: H3K4me1 is assumed to affect RNA Pol II processivity rather than transcription initiation/transition to elongation. Higher overall coverage of H3K4me1 across the locus therefore promotes distal termination of transcription by reducing the probability of early termination. H3K4me1 modifications are added co-transcriptionally, allowing distal termination to feedback onto further H3K4me1 addition. Conversely, proximal termination of both sense and antisense transcription leads to removal of H3K4me1, in line with the potential function of FLD as a demethylase, allowing feedback onto further proximal termination. Higher H3K27me3 coverage, rather than affecting processivity, is assumed to decrease the frequency of both sense and antisense transcription events (representing either transcription initiation or transition to productive elongation). The model describes constitutive deposition of H3K27me by Polycomb in the *FLC* nucleation region. H3K27me spreads to the rest of the locus through interactions with the nucleation region, through read-write feedback interactions. An experimentally determined gene loop mediating an interaction between the 5’ and 3’ ends of *FLC* is also incorporated in the model. Finally, in line with experimental evidence from mammalian systems and previously developed models for Polycomb, the model assumes that productive transcription promotes removal of H3K27me through both histone exchange and co-transcriptional demethylation, thereby implementing antagonism between transcription and Polycomb silencing. These aspects are integrated into a fully stochastic spatiotemporal simulation of the *FLC* locus, where transcription events (sense and antisense) and histone modification levels (H3K4me1 and H3K27me0/1/2/3) are simulation outputs. Full details of the model can be found in the Methods and Supplementary Information.

### FCA and FLD mediate a transcription-coupled repression mechanism that is sufficient to generate analogue regulation

Our previous analysis of the *fca* allelic series highlighted that differences in overall expression between genotypes appeared to have two components – a digital ON/OFF mode where we observed clear differences in the number of *FLC* OFF cells, and an analogue mode, with quantitatively different levels of *FLC* in the ON cells (Fig. 1B). Intermediate expression was associated with a slower switch off over time into the digital OFF state. Importantly, we also observed similar behaviour of *FLC* expression in single cells for a different intermediate mutant of *FCA*, called *fca-4* (Fig. S5C and (Antoniou-Kourounioti et al., 2023)), showing effects are not allele-specific, but related to the regulatory function of FCA at *FLC*. Overall, this demonstrates that FCA quantitatively modulates transcription and influences the timing of the switch into the digital OFF state.

We first tried to see if the model could capture the quantitative variation in *FLC* expression (analogue mode) in the absence of the Polycomb module (see model schematic in Fig. 3A). To represent the difference in FCA function between the three genotypes, we used a single parameter controlling the probability of FCA mediated proximal termination of transcription. We then simulated the model with different values for this parameter representing different levels of functionality of the FCA protein. We set the parameter to its highest value for L*er*, an intermediate value for *fca-3*, and lowest for *fca-1* (Fig. 3B). With these parameter settings we found that the model, with just transcription-coupled repression (Fig. 3A) was able to recapitulate the analogue mode of regulation, with the simulations showing the lowest frequency of productive transcription events for L*er*, highest for *fca-1*, and intermediate for *fca-3* (Fig. 3C). Consistent with the relationship between productive transcription and H3K4me1 in the model, the levels of this modification were also predicted to be lowest in L*er*, intermediate in *fca-3* and highest in *fca-1* (Fig. 3C). While the model simulation output shown in Fig. 3 is qualitatively consistent with our previous observations, note that the specific model parameter values used here are chosen to obtain a quantitative fit for the full model to experimentally measured histone modification levels as we explain below.

### Model predicts differences in the rate of Polycomb silencing establishment in the different genotypes

With the analogue module constrained in this way, we simulated the full model, including the Polycomb digital switching module (Fig. 3A). Importantly, this model recapitulated the experimentally observed reduction in *FLC* transcriptional output (Fig. 3D), as well as the digital switch to stable silencing (one-way switch) over time (Fig. 3E) observed using *FLC-*Venus imaging in root meristematic cells (Antoniou-Kourounioti et al., 2023). For L*er*, where productive transcription is lowest, the model predicted a rapid switch to the digital OFF/Polycomb silenced state (Fig. 3E, Fig. S4B(i)). For *fca-3*, the model predicted a slow switch to the digital OFF state, while even slower switching is predicted for *fca-1* (Fig. 3E, Fig. S4B(i)). In all cases, the switching was predicted to be one-way, with the Polycomb silenced state being highly stable.

Thus, the combined model of transcription-coupled repression and Polycomb silencing predicts that the level of productive transcriptional activity set by FCA/co-transcriptional silencing/FLD (analogue module) determines the level of transcriptional antagonism to H3K27me3. The level of transcriptional antagonism in turn dictates the rate of establishment of the Polycomb silenced state.

### Model predictions are consistent with observed chromatin changes over time

We next used the combined model to predict population levels of H3K4me1 and H3K27me3 at *FLC* in the three genotypes, predicting low H3K4me1 and high H3K27me3 levels in L*er*, intermediate levels of both modifications in *fca-3*, and high H3K4me1 and low H3K27me3 in *fca-1*. Starting from an active chromatin state with full H3K4me1 coverage, the model also predicted a progressive reduction in H3K4me1 over time in all genotypes, with a low steady state level quickly attained in *Ler*, and a slower reduction in *fca-3* and *fca-1*, with *fca-1* being the slowest (see red bars in Fig. 4D). In all genotypes, this reduction is combined with an accumulation of H3K27me3 as more loci switch to the Polycomb silenced state (see red bars in Fig. 4C).

**Figure 4:**
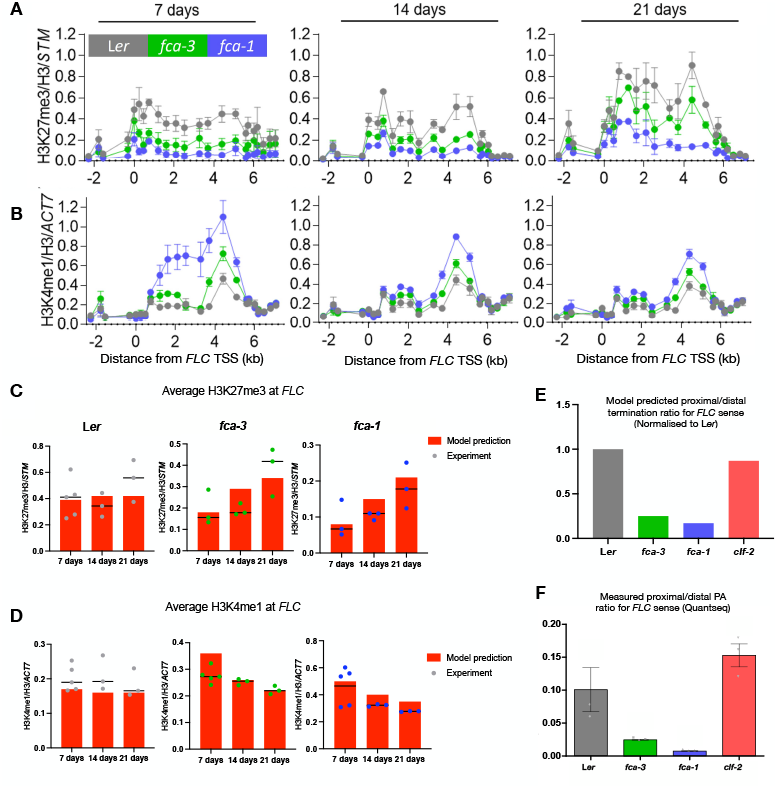
**(A**,**B)** ChIP-qPCR time course measurements of H3K27me3 (A), and H3K4me1 (B) across the *FLC* locus in the parental L*er* genotype and *fca* mutants. H3K27me3 levels are shown normalised to H3 and to control locus *STM*. H3K4me1 levels are shown normalised to H3 and to the control locus *ACT7*. Each dot represents an amplicon. Error bars in (A) represent mean +/- s.e.m. (n = 5 biological replicates only for L*er* at 7 days and n=3 biological replicates in all other cases). Error bars in (B) represent mean +/- s.e.m. (n = 5 biological replicates for L*er, fca-3*, and *fca-1* at 7 days and n=3 biological replicates in all other cases). **(C**,**D)** Experimental validation of model predictions. Vertical red bars indicate model predicted level. All model predicted histone modification levels are averaged over the locus and averaged over 1000 simulated copies of the locus started in an active state with full H3K4me1 coverage. Horizontal black bars indicate median of the biological replicates. **(C)** Comparison of model predicted H3K27me3 levels and changes over time in the different genotypes, with the measured levels shown in (A), averaged across 15 primers covering the *FLC* locus. **(D)** Similar comparison of model predicted H3K4me1 levels with the experimental data shown in (B). **(E**,**F)** Model predicted (E) and measured (Quantseq) (F) ratios of proximal to distal termination for *FLC* sense transcription, in the parental L*er* genotype, *fca* mutants and a *clf* mutant. Measured ratios were computed using the Quantseq data for 7 day old seedlings shown in Fig. 2B and Fig. S4C(i). Error bars in (F) represent mean +/- s.e.m. (n = 3 biological replicates). The genomic coordinates used to define the proximal and distal termination clusters are shown in the supplementary file FLC_polyA_cluster_coords.txt. The read counts within these clsuters for the different genotypes are provided in the supplementary file FLC_termination_clusters.txt. Model predicted ratios are normalised to the predicted ratio for L*er*. Model predicted ratios were computed at the 7 day timepoint (averaged over the last 0.25 days leading up to this timepoint) for all four genotypes (average over 1000 simulated copies started in an active state with full H3K4me1 coverage). The model qualitatively captures the relative changes in the proximal to distal ratio between mutants. Note that absolute predictions of the ratios would require estimates of lifetimes for the proximal and distal polyadenylated forms in each mutant.

To validate these predictions, we performed a time course measurement of silencing H3K27me3 and the activating modifications H3K4me1 and H3K36me3 over the *FLC* locus by chromatin immunoprecipitation followed by quantitative PCR (ChIP-qPCR), examining their levels in seedlings at 7, 14 and 21 days after sowing. For each genotype and timepoint, we use the Polycomb silenced *STM* locus as the positive control for H3K27me3 and the actively expressed *ACT7* locus as the positive control for H3K4me1 (conversely *STM* and ACT7 function as negative controls for the activating modifications and H3K27me3 respectively). See Fig. S4C(ii,iii,iv) for a comparison of the levels measured at *FLC* and these control loci at the 7 day timepoint. Consistent with model predictions, at all timepoints we see that *fca-3* shows intermediate levels of H3K4me1 and H3K27me3, as compared to *fca-1* and Ler, with primers across the *FLC* locus showing a consistent trend (Fig. 4A,B). This trend is reinforced by a statistical comparison between the three genotypes of the averaged H3K27me3 and H3K4me1 levels across the locus (Fig. S4A (i,ii)).

For validating the model using the timecourse ChIP data we tried to fit the model (adjusting parameters for the transcription-coupled repression mechanism) so that it captures the observed trends. The fitted parameters are indicated in the supplementary tables (Tables 1-4). We note that a direct quantitative comparison between the model predictions and the ChIP-qPCR data is difficult because the simplest model would assume that the switching dynamics happens at all loci, while the ChIP-qPCR analysis uses whole seedling tissue, including tissue where the switching may be inhibited by other means. To account for this possibility in the model, when comparing the model predictions to the ChIP-seq data, we include a subpopulation of ‘active’ *FLC* copies with full H3K4me1 coverage and no H3K27me3 coverage, to compute a corrected model predicted level of both modifications. The changes in histone modifications over time are then qualitatively consistent with the model (Fig. 4C,D). Experimental H3K27me3 levels rise in *fca-3* and *fca-1* (the increase is statistically significant in *fca-3* between 14 and 21 days, though narrowly not for *fca-1*: p=0.048, sig. threshold=0.0253 (Holm-Sidak corrected), Fig. S4A(iii)). H3K4me1 levels fall in *fca-3* and *fca-1* over time (statistically significant for both *fca-3* and *fca-1* between 14 and 21 days, Fig S4A(iv)) as more loci switch to the digital OFF state. In L*er*, H3K4me1 shows essentially no change over the experimental timepoints, while the H3K27me3 also does not show a statistically significant change on average across the locus (Fig. S4A(iii, iv)). The lack of significant change in experimentally measured H3K27me3 and H3K4me1 in L*er* are consistent with steady state levels having been reached by the 7-day timepoint. The model predicts faster dynamics of H3K4me1 removal (through more frequent proximal termination), and faster establishment of H3K27me3 in L*er*, so a rapid approach to steady state is consistent. The activating H3K36me3 also shows a similar pattern, with *fca-3* exhibiting intermediate levels, and the overall levels decreasing over time (Fig. S4B(iii)).

**Table 1:** Model parameter values: H3K27me.

| Parameter | Description | Value used in model | Details |
| --- | --- | --- | --- |
| $N$ | Number of histones | 60 | Approximately 6kb size of the *FLC* locus |
| $k_{me}$ | PRC2-mediated methylation rate (histone<sup>-1</sup> s<sup>-1</sup>) | $8 \cdot 10^{-6}$ | Parameter value fitted to genome-wide H3K27me3 dynamics in Berry et al., 2017 |
| $k_{me0-1}$ | PRC2-feedback mediated methylation rate (me0 to me1) (histone<sup>-1</sup> s<sup>-1</sup>) | $9k_{me}$ | Following the model in Berry et al., 2017 |
| $k_{me1-2}$ | PRC2-feedback mediated methylation rate (me1 to me2) (histone<sup>-1</sup> s<sup>-1</sup>) | $6k_{me}$ | Following the model in Berry et al., 2017 |
| $k_{me2-3}$ | PRC2-feedback mediated methylation rate (me2 to me3) (histone<sup>-1</sup> s<sup>-1</sup>) | $k_{me}$ | Following the model in Berry et al., 2017 |
| $\gamma_{me2-3}$ | Noisy methylation rate (me2 to me3) (histone<sup>-1</sup> s<sup>-1</sup>) | $k_{me}/20$ | Following the model in Berry et al., 2017 |
| $\gamma_{me1-2}$ | Noisy methylation rate (me1 to me2) (histone<sup>-1</sup> s<sup>-1</sup>) | $k_{me1-2}/20$ | Following the model in Berry et al., 2017 |
| $\gamma_{me0-1}$ | Noisy methylation rate (me0 to me1) (histone<sup>-1</sup> s<sup>-1</sup>) | $k_{me0-1}/20$ | Following the model in Berry et al., 2017 |
| $\gamma_{me2-3}^{NR}$ | De novo methylation rate (me2 to me3) in Nucleation region (histone<sup>-1</sup> s<sup>-1</sup>) | $0.35 \gamma_{me2-3}$ | Fitted in this study |
| $\gamma_{me1-2}^{NR}$ | De novo methylation rate (me1 to me2) in Nucleation region (histone<sup>-1</sup> s<sup>-1</sup>) | $0.35 \gamma_{me1-2}$ | Fitted in this study |
| $\gamma_{me0-1}^{NR}$ | De novo methylation rate (me0 to me1) in Nucleation region (histone<sup>-1</sup> s<sup>-1</sup>) | 0.35 $\gamma_{me0-1}$ | Fitted in this study |
| $\beta$ | Relative local PRC2-activity | 0.7 | Fitted in this study |
| $\rho_{me2}$ | Relative activation of PRC2 by H3K27me2 | 0.1 | Following the model in Berry et al., 2017 |
| $\gamma_{dem}$ | Noisy demethylation rate of H3K27 (histone<sup>-1</sup> s<sup>-1</sup>) | $8.8 \cdot 10^{-7}$ | Fitted in this study |

**Table 2:** Model parameter values: Nucleosome interaction parameters for PRC2 read-write feedback.

|  |  |  |  |
| --- | --- | --- | --- |
| $\xi_{NR}^{NR} = \xi_{LR}^{NR} = \xi_{NR}^{LR} = \xi_{LR}^{LR}$ | Interactions within nucleation and looping regions | 1 (*Ler*, *fca-3*, *fca-1*), 0.8 (*clf-2*, *fca-3* *clf-2*) | Fitted in this study |
| $\xi_{NR}^{Body} = \xi_{LR}^{Body}$ | Nucleation and looping region contributions to methylation of gene body nucleosomes | 0.5 (*Ler*, *fca-3*, *fca-1*)<br>0.05 (*clf-2*, *fca-3* *clf-2*) | Fitted in this study |
| $\xi_{Body}^{NR} = \xi_{Body}^{LR}$ | Gene body contributions to methylation of nucleation and looping region nucleosomes | 0.2 (*Ler*, *fca-3*, *fca-1*)<br>0.12 (*clf-2*, *fca-3* *clf-2*) | Fitted in this study |
| $\xi_{Body}^{Body}$ | Interactions within gene body region | 0.01 | Fitted in this study |

**Table 3:** Model parameter values: Transcription and PA site choice.

|  |  |  |  |
| --- | --- | --- | --- |
| $\alpha_{sense}$ | Trans-acting gene activation for sense transcription | 1 | Fixed to 1 for *FLC* sense transcription |
| $\alpha_{antisense}$ | Trans-acting gene activation for antisense transcription | 0.2 | Set to a lower value relative to $\alpha_{sense}$, consistent with lower frequency of antisense transcription compared to sense transcription under pre-cold warm conditions (Rosa et al., 2016) |
| $f_{min}$ | Minimum transcription initiation | $f_{max}/60$ | Fitted in this study |

| | rate ($s^{-1}$) | | |
| --- | --- | --- | --- |
| $f_{max}$ | Maximum transcription initiation rate at *FLC* ($s^{-1}$) | $0.58 \cdot 10^{-3}$ | *FLC* sense transcription initiation frequency computed using the same approach as Ietswaart et al., 2017, but using single cell mRNA counts and *FLC* mRNA half-life measured for *fca-1* in Antoniou-Kourounioti et al., 2023 |
| $P_T$ | H3K27me3 coverage threshold for full silencing of transcription | 1/3 | Following the model in Berry et al., 2017 |
| $\delta_{proximal}$ | Residual probability of proximal termination | 0.05 | Fixed in this study |
| $\mu_{Distal}$ | Residual probability of Distal termination | 0.05 | Fixed in this study |
| $p_0$ | Parameter representing FCA independent proximal termination | 0.8 | Fitted in this study |
| $p_{FCA}$ | Parameter representing FCA mediated proximal termination | 0.2 (*Ler*, *clf-2*), 0.15 (*fca-3*), 0 (*fca-1*) | Fitted in this study |
| $\eta$ | Weight parameter for relative contribution of NR+LR H3K27me3 to transcriptional silencing | 0.25 | Fitted in this study |

**Table 4:** Model parameter values: Transcription effects on modifications.

|  |  |  |  |
| --- | --- | --- | --- |
| $p_{dem}$ | Demethylation probability of H3K27me (histone$^{-1}$ distal transcription$^{-1}$) | 0.1 | Fitted in this study |
| $p_{ex}$ | Histone exchange probability (histone$^{-1}$ distal transcription$^{-1}$) | $1 \cdot 10^{-3}$ | Following the model in Berry et al., 2017 |
| $p_{dem}^{FLD}$ | Demethylation probability of H3K4me1 (histone$^{-1}$ proximal transcription$^{-1}$) | 0.5 | Fitted in this study |
| $p_{K4me}^{Transc}$ | Methylation probability for unmodified H3K4 during a distally terminated transcription event | $100 \frac{k_{me}}{f_{max}}$ | Computed as 100 times the PRC2-mediated methylation rate $k_{me}$ (histone $^{-1}$ s $^{-1}$ ), then normalised by transcription frequency ( s $^{-1}$ ) to obtain probability per histone per transcription event. |

Overall, the combined model of transcription-coupled repression – mediated by proximal termination of transcription, and Polycomb silencing, successfully predicts both the relative levels of the modifications between genotypes and how these change over time.

### Model predicts relative differences in proximal termination between genotypes

To validate the model predicted *FCA* dependent differences in proximal termination, we compared these predictions to our Quant-seq data (data shown in Fig. 2B). We compared the predicted ratio of proximal to distal termination events to the measured PA reads. To compute this ratio for the Quant-seq data, we defined a Proximal termination cluster covering a range of termination sites close to the 5’ end of the gene. Our definition of the proximal termination cluster is based on the proximal termination of *FLC* sense transcripts we have previously reported (Schon et al., 2021). The genomic coordinates are provided in a supplementary text file: FLC_termination_clusters.txt. A quantitative fit of these ratios is not attempted here, as we do not know turnover rates of the different transcripts. However, the predicted trends in the differences in the ratio of proximal to distal polyadenylation of sense transcripts between the *FCA* mutants and the parental genotype qualitatively matched the trend for the measured ratios of *FLC* sense transcription in our Quant-seq dataset (Fig. 4E,F; Fig. S4C(i), see Fig. S4D for individual replicates).

### Disrupting a Polycomb methyltransferase compromises the one-way switch to a Polycomb OFF state but does not disrupt FCA mediated regulation

The model predicts that the transcription coupled repression mechanism is sufficient to generate the analogue repression and that this repression controls the antagonism of transcription to Polycomb, thus enabling the switch to Polycomb digital silencing. We tested these predictions using a Polycomb mutant – a loss of function mutant of *CLF*. This methyltransferase is known to be required for spreading of H3K27me3 from Polycomb nucleation peaks at many Arabidopsis targets (Shu et al., 2019), and specifically at *FLC*, both in developmental (Shu et al., 2019) and cold induced silencing contexts (Yang et al., 2017). This mutant is not expected to completely disrupt the ON/OFF silencing mode, as H3K27me3 nucleation alone, which still occurs in *clf*, gives a metastable Polycomb silenced state (Yang et al., 2017). However, only after the H3K27me3 spreads across the whole locus is the Polycomb silenced state fully stable in rapidly dividing cells (Yang et al., 2017). Thus, the maintenance of the OFF state is expected to be compromised in *clf* (Fig. 5A, B). Therefore, in a *clf* mutant background, the model makes two predictions: (i) that the continued slow reduction in overall *FLC* expression seen in L*er, fca-3*, and *fca-1*, i.e., an overall reduction in the number of actively transcribed *FLC* copies – which is a consequence of a fully stable Polycomb digital switch – will be missing in a *clf* mutant background, due to leakage out of the metastable Polycomb OFF state (Fig. 5A,B); (ii) that the analogue differences generated by *fca* mutants will persist, since the FCA mediated repression can function independently of H3K27me3.

**Figure 5:**
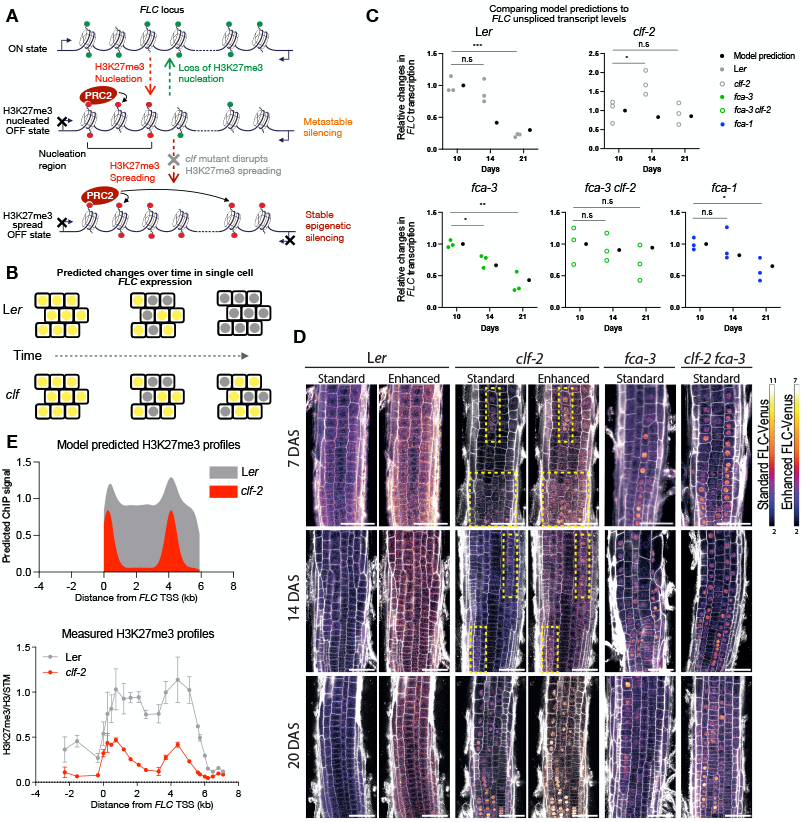
**(A)** Schematic showing the rationale for examining the *clf* mutant. Transition from a Polycomb nucleated state to a spread state is prevented in a *clf* mutant. The lack of stability of a nucleated state allows switching back to a digital ON state. Green circles: active modifications (H3K4me1); red circles: repressive modifications (H3K27me3). **(B)** Consequently, the reduction over time in the number of ON cells observed in the parental L*er* genotype is expected to be disrupted in the *clf* mutant. **(C)** Comparison of model predicted changes in *FLC* transcription, with whole seedling *FLC* unspliced transcript levels measured by qPCR in the *fca* genotypes (L*er, fca-3, fca-1*) and the *clf* mutants *(clf-2* and *fca-3 clf-2* double mutant). Experimental data is normalised to measured levels for *UBQ10*. Statistical comparisons for the experimental data were carried out using the Student’s t-test and corrected for multiple comparisons using the Holm-Sidak approach (*, **, *** indicate significance thresholds of 0.05, 0.01, and 0.001 respectively). Model predictions correspond to the predicted frequency of *FLC* sense distal transcription events at the corresponding timepoints, averaged over 1000 simulated loci. Simulations for L*er, fca-3, fca-1* are identical to those used to predict changes in histone modification levels (Fig. 4). Simulations for the *clf* mutants *(clf-2, fca-3 clf-2*) use changed parameter values for reactions involving H3K27me3 addition by PRC2 – see Supplementary Information and Supplementary Table 2. For comparison of relative changes, both model predictions and data are normalised to the level at 10 days. **(D)** FLC-Venus time-course imaging in the different genetic backgrounds. Representative confocal images of FLC-Venus signal at three timepoints in the epidermis of the root meristem of the parental genotype Ler, the single mutants *clf-2*, and *fca-3*, and the double mutant *fca-3 clf-2*. Grey shows the cell wall dye propidium iodide; FLC-Venus intensity is indicated by colour maps for standard and enhanced images, respectively. The same settings were used for imaging of all time points. The same image presentation is used for standard and enhanced images. Brightness and contrast of FLC-Venus signal were increased to generate the enhanced images (Note different scale on colour maps). Yellow boxes in *clf-2* highlight ON cell files at DAS7/14. Scale bar, 50 □ μm. DAS stands for days after sowing. **(E)** The model can capture the differences between L*er* and *clf-2*, in the spatial profile of H3K27me3 across the *FLC* locus. Top panel: model predicted profiles of H3K27me3 coverage (see Supplementary Information for detailed description of simulation). Model predicted profile is steady-state average over 100 simulated loci, started in an active chromatin state with full H3K4me1 and equilibrated over a time duration of 10 cell cycles. Bottom panel: H3K27me3 profiles measured by ChIP-qPCR in 14-day old seedlings. Error bars represent mean +/- s.e.m. (n = 3 biological replicates). Each dot represents an amplicon.

To validate this prediction, we measured *FLC* expression (measuring spliced and unspliced *FLC* transcripts by qPCR) at the whole plant level, over the same developmental time course in five different genotypes (Fig. 5C shows unspliced *FLC;* Fig. S5A shows spliced *FLC)*: in the parental L*er, clf-2* (Goodrich et al., 1997), *fca-3*, the *fca-3 clf-2* double mutant, and *fca-1*. We compared these data to predicted *FLC* sense transcriptional activity from simulations of the full model for L*er, fca-3* and *fca-1*, as well as for L*er* and *fca-3* with the PRC2 read-write feedback parameters changed to capture a *clf* mutant. We did not attempt to fit to absolute levels of *FLC* transcripts here, since the measured levels also depend on RNA stability, which we do not measure experimentally or include in our model. Therefore, we compare changes relative to the measured levels at the first (10 day) timepoint. We saw that, as predicted by the model, the slow reduction in expression is compromised in the *clf-2* and *fca-3 clf-2* mutants (Fig. 5C, Fig. S5A) – neither of these mutants exhibits a statistically significant *reduction* in *FLC* expression over the timecourse. We note that *clf-2* exhibits an increase at the 14 day timepoint, (statistically significant only for unspliced *FLC*) that the model cannot capture.

Also consistent with predictions for the *clf* mutants, we observe that the analogue differences associated with FCA are maintained, i.e., the expression level in the *clf-2* mutant is clearly lower than in *fca-3* at all timepoints (Fig. S5B), indicating that the FCA mediated transcription coupled repression is the main factor underlying analogue differences. We also compared the model predicted changes in *FLC* transcriptional output (i.e., frequency of distally terminated sense transcription events) for *Ler, fca-3*, and *fca-1* to the above data (Fig. 5C, Fig. S5A). Consistent with previous data (Antoniou-Kourounioti et al., 2023), and the observed chromatin changes (Fig. 4), all three genotypes *Ler, fca-3*, and *fca-1* exhibit statistically significant changes in *FLC* expression over the time course (both spliced and unspliced *FLC*). The relative changes predicted by the model are overall quantitatively consistent with the measured changes in both unspliced (Fig. 5C) and spliced *FLC* (Fig. S5A) (note that the model predictions are identical for spliced and unspliced – the model does not distinguish between the two). We note that in some cases, between 10 days and 14 days, the model predicted changes are larger than the measured changes, notably, for unspliced *FLC* in L*er* (Fig.5C) and spliced *FLC* in *fca-3* and *fca-1* (Fig. S5A). However, since the model was only parametrised to fit the ChIP time course data (Fig. 4), the overall satisfactory quantitative fit observed for *FLC* expression with the same parameter values provides further support for the validity of the model.

In addition to *FLC* expression at the whole plant level, we also conducted time course imaging of Venus tagged FLC in the roots in the *clf* mutants (Fig. 5D), following the same approach used to analyse the *fca* alleles in (Antoniou-Kourounioti et al., 2023). Again, as predicted, *clf-2* showed clear differences to the L*er* genotype, with ON cells being visible at all three timepoints. This is consistent with *FLC* switching both ON to OFF and OFF to ON as described above, and the observed lack of reduction in *FLC* expression at the whole plant level (Fig. 5C). We note that both *clf-2* and *fca-3 clf-2* exhibited large variability between roots, with some roots showing essentially no *FLC* ON cells (Fig. S5C). This is possibly due to the widespread misregulation of growth and developmental genes resulting from the compromised Polycomb silencing in this mutant. Nevertheless, consistent with the model predictions, we see that the slow Polycomb digital switch to OFF is disrupted in *fca-3 clf-2*: when comparing roots exhibiting ON cells in the double mutant to *fca-3* roots, the double mutant overall shows more ON cells at later timepoints.

The model also predicts a similar, high ratio of proximal to distal termination in the *clf* mutant and the L*er* genotype (Fig. 4E), even though overall transcription is lower in L*er*. This is because the FCA/FLD mediated transcription coupled mechanism is not disrupted in the *clf* mutant model, so that the promotion of proximal termination by this mechanism, and the resulting feedback, remain intact. We measured the ratio of proximal to distal termination of the *FLC* sense transcript in the *clf* mutant, using our strand-specific Quant-seq data (Fig. S4C (i)). Consistent with the model prediction, it exhibits a similar ratio to the L*er* genotype (Fig. 4F).

As further validation of the model, we then compared the predicted H3K27me3 levels at the locus in a *clf* mutant to experimentally measured levels. The lack of stability of the nucleated state in the model means that it predicts an H3K27me3 peak, but at significantly lower levels relative to a spread state. The incorporation of the *FLC* gene loop in the model, as outlined above, causes the model to generate a second peak in the *FLC* 3’ region, in the region that interacts strongly with the nucleation region. Consistent with these predictions, we find that the ChIP profile of H3K27me3 in *clf-2* exhibits two peaks, at significantly lower levels relative to the spread state in L*er* (Fig. 5E). Our results also aligned well with previously published genome-wide H3K27me3 data for a *clf* mutant in a Col-0 background (Shu et al., 2019). Overall, predictions from the model for *clf* mutants match well with our experimental data.

### The repressor FLD is co-transcriptionally targeted to *FLC* but requires FCA mediated proximal termination for its repressive function

We next investigated the mechanism of FLD functionality in the co-transcriptional repression mechanism. In apparent contradiction to its role as a repressor, in previously reported data, FLD occupancy was found to be significantly higher in a high transcriptional state of *FLC*, relative to a Polycomb silenced low transcriptional state. Specifically, high FLD levels were assayed by ChIP-seq of a transgenic *fld-4; 3xFLAG-FLD* in the Col*FRI* background, where the *FLC* activator FRIGIDA sets up a high transcriptional state (Inagaki et al., 2021). In contrast, in a Col-0 background, where FRI is not functional, *FLC* is Polycomb silenced and FLD shows low enrichment.

The model allows us to explain this counter-intuitive behaviour of FLD. Since FLD function is coupled to transcription and termination in the model, FLD targeting must be co-transcriptional, consistent with the molecular and genetic analysis in the accompanying paper (Mateo-Bonmatí, 2023). FLD levels at the locus must therefore be low in a Polycomb silenced state, where transcription events are infrequent. Such co-transcriptional FLD targeting is consistent with the genome-wide correlation between FLD enrichment and transcription as measured by Pol II ChIP and transcript amounts in (Inagaki et al., 2021). Furthermore, by capturing the digital switch off in L*er* and *fca-3*, the model predicts that FLD enrichment would reduce over time in these genotypes. In essence, FLD is only needed transiently to generate a lowly-transcribed state capable of nucleating Polycomb silencing, as transcription antagonises Polycomb. Following Polycomb nucleation and spreading of H3K27me3, FLD recruitment to the locus is reduced due to the lack of transcription.

The experimental validation of the model assumption of FLD targeting coupled to transcription termination, is presented in the accompanying paper (Mateo-Bonmatí, 2023), where we elucidate the links between the FLD complex (Fang et al., 2019) and CPF components. By ChIP-qPCR analysis of FLD enrichment at *FLC*, comparing high transcription states induced by two very different genetic backgrounds (*fca-9* and *FRI*) with a Polycomb silenced state (Col-0), we show that high FLD enrichment at *FLC* accompanies high transcription, irrespective of FCA function. Our model also supports the scenario proposed by Inagaki et al., (2021) in which FLD downregulates histone H3K4me1 around termination sites in regions with convergent overlapping transcription genome-wide in Arabidopsis.

We also tested whether FLD enrichment would change over time, as expected from the model predicted switch to the Polycomb silenced state. We measured 3xFLAG-FLD enrichment by ChIP-qPCR over a time course in a Col-0 background (as high-expressing *fca-9* and FRI are not expected to exhibit a digital switch OFF). Consistent with model predictions, we observed a reduction over time in FLD enrichment across *FLC* (Fig. 6B), mirroring the reduction in *FLC* transcription levels over time also observed in Col-0 by qPCR (Fig. 6A). Levels at the initial 7-day timepoint were high and declined to very low levels (almost background) by 21 days. Consistently, both 7-day and 14-day average FLD levels across the locus showed statistically significant differences to the control sample, while the 21-day levels did not (Fig. S6).

**Figure 6:**
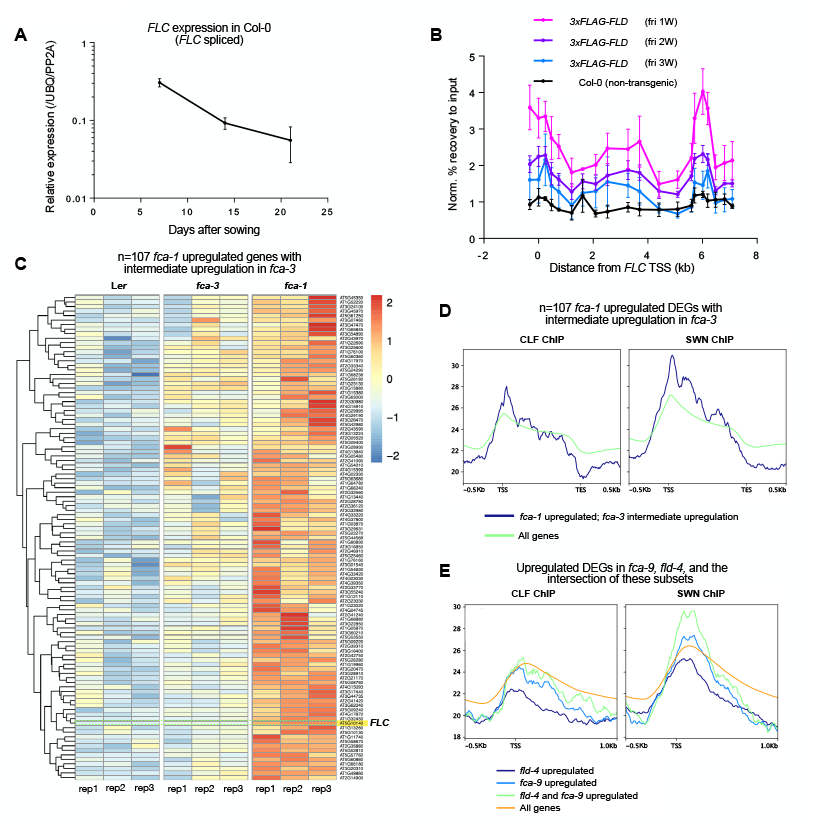
**(A)** Changes over time in spliced *FLC* expression in Col-0 seedlings, measured by time-course qPCR. Error bars represent mean +/- s.e.m. (n = 3 biological replicates). Data is shown normalised to the measured levels for *UBQ10* and *PP2A*. **(B)** Changes over time in FLD targeting to the *FLC* locus, measured by time-course ChIP-qPCR quantification of 3xFLAG-FLD in seedlings (Col-0 background) (normalized to IGN5). Error bars represent mean +/- s.e.m. (n = 3 biological replicates). **(C)** Heatmap generated from analysing genome wide Quant-seq data, showing changes in transcriptional output at the 107 targets that are significantly upregulated in *fca-1* and also show intermediate (but not necessarily statistically significant) upregulation in *fca-3*. **(D)** Metagene plots showing enrichment of the PRC2 methyltransferases SWN and CLF at the 107 targets (Differentially Expressed Genes, DEGs) shown in (C), compared to their enrichment genome-wide. **(E)** Metagene plots comparing SWN and CLF enrichment at *fca-9* upregulated, *fld-4* upregulated, and overlapping DEGs, with their genome-wide enrichment.

Together, these observations are consistent with co-transcriptional FLD targeting to the *FLC* locus, with its repressive activity dependent on FCA-mediated proximal polyadenylation of sense and antisense transcription. In genotypes such as *fca-9*, where FCA is non-functional, or in the presence of the transcriptional activator FRIGIDA, which antagonises proximal PA by FCA (Schon et al., 2021), FLD continues to be co-transcriptionally targeted, but fails to carry out its repressive function. This highlights an important aspect relating genome-wide localisation of a regulator to its function: factors that are found to localise at active transcription sites may not just be transcriptional activators. As is the case for FLD, such factors could be essential components for shutting down transcription in specific developmental contexts.

### FCA mediated transcription-coupled repression potentially controls many Polycomb targets

We next analysed our genome-wide Quant-seq data to examine whether the FCA-mediated transcription coupled mechanism could be applicable to a wider range of targets beyond *FLC*, and whether this mechanism is also linked to Polycomb silencing at other targets. We first analysed differential upregulation in *fca-1* (null) relative to L*er*, which showed that 130 targets were significantly upregulated in *fca-1*. Of these 130 targets, 107 were upregulated to an intermediate level in *fca-3* (without however showing statistically significant upregulation in *fca-3*, probably because of the intermediate nature of this genotype), hence behaving similar to *FLC* (Fig. 6C). We then examined whether these 107 targets were also Polycomb targets. We used published ChIP-seq data for the Arabidopsis PRC2 methyltransferases CLF and SWN (Shu et al., 2019) in the Col-0 genotype, and generated a metagene plot for CLF and SWN enrichment across the 107 targets, comparing it to the whole genome metaplot for CLF and SWN enrichment. This showed a clear higher enrichment of these Polycomb components at these 107 putative FCA targets, particularly close to the Transcription Start Site (TSS), compared to the genome-wide average (Fig. 6D).

In order to examine whether FCA and FLD function overlap in general, we used genome-wide Quant-seq data in the null mutant genotypes *fca-9* and *fld-4* and the corresponding parental genotype Col-0, to carry out a similar analysis. We found that 91 genes were significantly upregulated in *fca-9* compared to the parental genotype, Col-0. We first examined whether these targets overlap with those observed in *fca-*1. Our data shows extensive differential gene expression between the Col-0 and L*er* parental genotypes (not shown), and these differences are reflected in the respective mutants in these backgrounds. It is therefore not surprising that we find a relatively small overlap of 10 targets between the *fca-1* upregulated set and the *fca-9* upregulated set, even though these are both null mutants of *FCA*.

Analysis of *fld-4* relative to Col-0 showed a larger number of upregulated targets (193), consistent with the proposed wider role for FLD in coordinating convergent transcription genome-wide (Inagaki et al., 2021). We found that 48 targets were significantly upregulated in both *fca-9* and *fld-4*, indicating that these factors may be working together at a number of genes. As above, we compared metagene plots of CLF and SWN enrichment across these 48 targets, as well as for the *fca-9* and *fld-4* singly upregulated subsets, in a whole genome metagene plot for CLF and SWN enrichment. For the *fca-9* upregulated genes, we saw slightly higher enrichment of SWN but not of CLF around the TSS compared to the genome-wide average (Fig. 6E). Interestingly, for the *fld-4* upregulated subset, we observe lower enrichment of both SWN and CLF compared to the genome-wide average (Fig. 6E). This is consistent with the co-transcriptional targeting of FLD, but where the majority of the targets do not couple to Polycomb. However, the metagene plot for the 48 overlapping targets compared to the genome wide-average shows a further increase in SWN enrichment (and a smaller increase for CLF) around the TSS than seen for the *fca-9* upregulated targets alone (Fig. 6E). This suggests that FLD function may be linked to silencing by PRC2 in specific contexts, where FLD action is facilitated by FCA mediated proximal polyadenylation.

Together, these analyses indicate that FCA/FLD mediated transcription-coupled repression may be working at a broad range of targets, and that at these targets, as at *FLC*, this repression works in concert with the Polycomb system. We also note that multiple genes of general interest are present in the *fca-1* and *fca-9/fld-4* upregulated lists, including: a Trithorax homologue (ATX1) implicated in development and multiple stress responses, genes such as GRP5 (Mangeon et al., 2010) and GASA9 (Yang et al., 2008) showing highly cell type specific expression, and several stress inducible genes (see Supplementary Table S2 for gene names of selected targets), all features that are consistent with Polycomb regulation.

## Discussion

How co-transcriptional processing mechanistically influences transcriptional output, and how this is linked to epigenetically stable Polycomb silencing is still not understood in any system. We have exploited the Arabidopsis *FLC* system to investigate these questions. We had shown that a transcription-coupled repression mechanism; specifically involving proximal transcription polyadenylation/termination mediated by the RNA-binding protein FCA interacting with 3’ processing machinery, can generate graded (analogue) transcriptional output. We had also shown that this mechanism was coupled to histone demethylase (FLD) activity. In the accompanying paper (Mateo-Bonmatí, 2023), we showed that APRF1, a structural component of a CPF-like phosphatase complex, directly links transcription termination with the FLD histone demethylase activity to alter the local chromatin environment. But how FLD activity can affect proximal termination, how this mechanism can generate analogue transcriptional output, and how this transcription-coupled repression links to a digital ON to OFF switch to PRC2 silencing remained obscure.

We built on our previous using the computational model described here, and used it to reveal how the level of productive transcriptional activity set by the transcription-coupled mechanism (FCA/FLD) determines the level of transcriptional antagonism to Polycomb silencing. The different levels of transcriptional activity set by the transcription coupled repression can dictate the timescale for establishment of Polycomb silencing (Fig. 7). We validated the model by a variety of means, including gene expression, histone modification and FLC-Venus imaging time courses in the wild-type and in cases disrupted via a Polycomb mutation. Our model also resolves the apparent contradiction between the observed FLD targeting to a transcriptionally active *FLC* locus, and its role as a repressor: FLD’s co-transcriptional recruitment, needed to lower transcriptional activity to pave the way for Polycomb silencing, is eliminated once full Polycomb silencing is achieved. Hence, factors localising at activated genes can in some instances be paradoxically essential for shutting down transcription.

**Figure 7:**
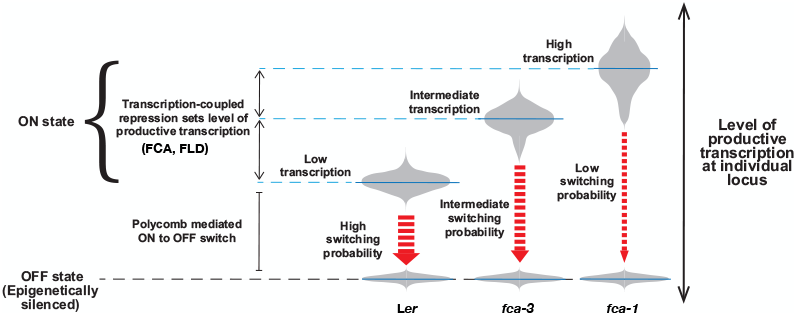
Schematic showing how analogue and digital regulation can combine in the context of Polycomb mediated silencing. The differences in thickness of the red arrows indicate the different rates of switching to the Polycomb silenced OFF state.

A key methodology to dissect the underlying mechanism was the construction of a computational model. The model was used as a tool to predict the levels of dynamically changing transcriptional and histone modification levels and to gain intuition about the underlying mechanisms. We chose to keep the model simple and interpretable but the model was still able to reproduce all the statistically significant trends in the data.

The transcriptional antagonism to Polycomb silencing, a fundamental aspect of the silencing mechanism, is also important in the mammalian and Drosophila contexts (Klose et al., 2013), (Holoch et al., 2021). In the plant context, this antagonism sets the timescale for the slow switch into the digital Polycomb silenced state, in an interlinked three-way association between chromatin, transcription and co-transcriptional processing (Graphical Abstract). Importantly, our model predicts slow/inefficient establishment of Polycomb silencing in the *fca-1* mutant. As a result, if silencing can be established in *fca-1* (independently of FCA), for example mediated by cold-specific PRC2 accessories in the case of cold-induced silencing, we predict that both silenced and active states can each be stably maintained. Hence, our conceptual framework also explains why cold-induced Polycomb silencing can be maintained in *fca* (or *fld*) mutants. The model predicts that these proteins are needed only for Polycomb silencing establishment during development, but not for its maintenance. In future work, it would be very interesting to specifically test this prediction by removing FCA and/or FLD at a later developmental stage after silencing is established.

The putative H3K4 demethylase FLD is known to be a key component of the FCA mediated transcription-coupled repression mechanism, indicating an important role for removal of H3K4me associated with and downstream of proximal termination. A key mechanistic feature of the model is the existence of a link also in the opposite direction: H3K4me1 levels influencing the processivity of RNA Pol II – with low H3K4me1 levels reducing processivity. Thus, the model contains a positive feedback loop (mutual inhibition) between proximal termination and H3K4me1. While chromatin modifications have been mechanistically linked to co-transcriptional splicing (Leung et al., 2019), links to Pol II processivity are only beginning to be explored, as seen from recent findings on Pol II regulation by the Integrator complex in mammalian systems (Stein et al., 2022; Wang et al., 2023). These links could also involve an effect on Pol II speed which can influence termination (Cortazar et al., 2019).

Our model also explains how the *FLC* transcriptional activator FRIGIDA can set up an active transcriptional state, preventing Polycomb silencing by functioning as an anti-terminator. Anti-termination factors that prevent the use of early termination sites have been implicated in tuning transcriptional output from eukaryotic polymerases (Gregersen et al., 2019). In the case of *FLC*, the transcriptional activator FRIGIDA has recently been shown to function at least partly by antagonising FCA mediated proximal cleavage and polyadenylation of *FLC* sense transcription at specific stages of embryo development (Schon et al., 2021). The presence of active FRIGIDA promotes distal polyadenylation and prevents *FLC* silencing, further supporting the view that FCA mediated proximal termination is essential for repression. Interestingly, FRIGIDA expression outside of a specific developmental stage fails to activate *FLC* expression (Schon et al., 2021): this reinforces the view that once Polycomb epigenetic silencing has been established, reactivating transcription is difficult, with simple upregulation of an activator being insufficient to overcome robust digital silencing. The model predicts that the locus is bistable in the absence of proximal-termination mediated repression, so that both active and Polycomb silenced states can be stably maintained (Fig. S4B(ii)). This is consistent with the FCA mediated mechanism not having a role in maintaining the Polycomb silenced state, as evidenced by the stable maintenance of cold-induced Polycomb silencing at *FLC* in both *fca* mutants and genotypes with active FRIGIDA.

Overall, we have revealed the mechanism of co-transcriptional silencing at *FLC*, an essential precursor for slow switching into the Polycomb silenced state. These features are included in an animation to help explain these molecular feedbacks, and how this mechanism can lead to expression of *FLC* in one of two stable expression states (supplementary file: Movie_1.mp4). It is particularly intriguing that many elements of this pathway, such as FLD and FCA, are conserved transcriptional/splicing regulators, that likely have functions at many more targets. Their core role in an integrated sense/anti-sense transcriptional circuitry likely singles out *FLC* as being acutely sensitive to their action. Indeed, our analysis of genome-wide Quant-seq data and overlap with ChIP-seq data for Polycomb components indicates that this transcription-coupled mechanism targets many genes and potentially works in concert with Polycomb at these targets. We therefore anticipate these mechanisms will have even broader relevance, with additional examples of such mechanisms emerging over time (Chen et al., 2023). Further investigating the links between co-transcriptional mechanisms and Polycomb silencing will therefore be an important focus for future research.

## Methods

### Plant materials

All the plants were homozygous for the indicated genotype. Seeds were surface sterilized in 40 % v/v commercial bleach for 10 min and rinsed 4 times with sterile distilled water. Seeds were then sown on standard half-strength Murashige and Skoog (MS) medium (0.22% MS, 8% plant agar) media plates and kept at 4°C in darkness for 3 days before being transferred to long day photoperiod conditions (16 h of light, 8 h dark).

### Plant growth (Imaging)

Seeds were surface-sterilized with 5% (v/v) sodium hypochlorite for 5 min, rinsed with sterile water 4x for 1 min and stratified at 4°C for 2 days in water and in the dark. Seeds were plated on MES buffered Murashige and Skoog (GM-MS) media, pH 5.8, containing respective antibiotics and grown on vertically oriented petri plates for 7 days in long-day conditions (16 h light/8 h dark, 22/20°C).

### Imaging

Time course imaging of FLC-Venus protein levels in the epidermis of L*er, fca* allelic mutants, the *clf-2* mutant as well as the *fca-3 clf-2* mutant, was performed using a Leica confocal Stellaris 8 microscope with a 20x multi-immersion objective (0.75 NA). Root tips (2 cm) were cut off and immersed in 1.5 μg/mL Propidium Iodide (PI, Sigma–Aldrich, P4864) in 1x PBS before being imaged immediately. The OPSL 514 laser was used at 5% power with 514 nm excitation for FLC-Venus and PI. In photon counting mode, Venus was detected between 518-550 nm with the HyD SMD2 detector, PI was detected between ∼600-650 nm. Images were acquired with a laser speed of 400 Hz, line accumulation of 6 (pixel dwell time of 2.43 μs), a Z-step size of 0.95 μm and a pinhole size of 1 AU. The same settings were used at all imaging time points to allow direct comparison between genotypes and time points. The representative images were projected using a single middle slice from the PI channel to show the cell outline, and 10 slices of FLC-Venus channel were maximum intensity projected. In standard image presentation, the dynamic range of the FLC-Venus channel was pushed from 0-255 to 2-11, in enhanced image presentation, the dynamic ranged was pushed from 2-7. In Supplementary Figure S5C, the dynamic range of FLC-Venus channel was enhanced from 0-255 to 2-20 for all images.

### RNA extraction and RT-qPCR

Total RNA was extracted as previously described (Box et al. 2011). Genomic DNA was digested with TURBO DNA-free (Ambion Turbo Dnase kit AM1907) according to manufacturer′s guidelines, before reverse transcription (RT) was performed. The RT reactions were performed with SuperScript IV reverse transcriptase (ThermoFisher, 18090010) following the manufacturer′s instructions using gene-specific primers. Primers are listed in Supplemental Table S1. For RNA sequencing experiments, RNA was further purified with the Qiagen Rneasy miniprep kit (74106).

### RNA sequencing

Library preparation and sequencing was performed by Lexogen GmbH. RNA integrity was assessed on a Fragment Analyzer System using the DNF-471 RNA Kit (15 nt) (Agilent). Sequencing-ready libraries were generated from 100ng of input RNA using a QuantSeq 3’ mRNA-Seq Library Prep Kit REV for Illumina (015UG009V0271) following standard procedures. Indexed libraries were assessed for quality on a Fragment Analyzer device (Agilent), using the HS-DNA assay and quantified using a Qubit dsDNA HS assay (Thermo Fisher). A sequencing-ready pool of indexed libraries were sequenced on an Illumina NextSeq 2000 with a 100-cycle cartridge using the Custom Sequencing Primer (CSP) at Lexogen GmbH.

### Bioinformatics analysis

Data analysis of RNAseq data was performed by Lexogen GmbH using their standard pipeline. Read Quality was assessed by FastQC and trimmed for quality and adapter content with cutadapt version 1.18 (Martin, 2011). Clean reads were mapped to the Arabidopsis reference genome (TAIR10) with STAR version 2.6.1a (Dobin et al., 2013). Raw reads have been deposited on Short Read Archive (SRA) under the reference PRJNA980626, and PRJNA1076161. To calculate proximal to distal count ratios, read ends were counted for each identified polyA cluster ((Schon et al., 2021), FLC_polyA_cluster_coords.txt). Proximal polyadenylation clusters are defined as termination sites within the first Exon and Intron of *FLC* while distal clusters are in close proximity to the canonical polyadenylation site. For each biological sample, ratios were calculated from the summed proximal and distal cluster counts (FLC_termination_clusters.txt).

### ChIP timecourse for histone modifications

2.5 g of 7-, 14-, or 21-day-old seedlings were crosslinked in 1% formaldehyde and grinded to fine powder. Nuclei were extracted using Honda buffer as described previously in Sun et al. 2013. In all histone ChIP reactions, sonication, immunoprecipitation, DNA recovery and purification were performed as previously described in Wu et al. 2016. The antibodies used were anti-H3 (ab176842), anti-H3K27me3 (Merck 07-449 and ab192985), anti-H3K36me3 (ab9050), anti-H3K4me1 (ab8895). All ChIP data were quantified by qPCR with primers listed in Table S1. Values were normalized to H3 and internal control to either STM or ACT7.

### H3K27me3 ChIP in *clf* mutant

2.0 g of 14-day-old seedlings were crosslinked in 1% formaldehyde and grinded to fine powder. Nuclei were extracted using Honda buffer as described previously in Sun et al. 2013. In all histone ChIP reactions, sonication, immunoprecipitation, DNA recovery and purification were performed as previously described in Wu et al. 2016. The antibodies used were anti-H3 (ab176842), and anti-H3K27me3 (ab192985). All ChIP data were quantified by qPCR with primers listed in Table S1. Values were normalized to H3 and internal control *STM*.

### FLD ChIP

2.5 g seedlings were crosslinked in 0.685 mg/mL of EGS [ethylene glycol bis(succinimidyl succinate)] and 1% formaldehyde and grinded to fine powder. Nuclei were extracted using Honda buffer as described for histone ChIP. Nuclei pellets were resuspended in 600 μl of RIPA buffer (1% NP-40, 0.5% Sodium Deoxycholate, 0.1 % SDS in PBS) and sonicated 5 times for 5 min (30s ON/ 30s OFF) at high mode in a Bioruptor Pico (Diagenode). IP were performed overnight at 4°C using 1.5 mg of pre-coated epoxy Epoxy Dynabeads (M-270) with 1.5 μl of anti-FLAG (Sigma, F1804) per reaction. Beads were then washed for 5 min twice with Low Salt Buffer (150 mM NaCl, 0.1% SDS, 1% Triton X-100, 2 mM EDTA, 20 mM Tris-HCl), twice with Hight Salt Buffer (500 mM NaCl, 0.1% SDS, 1% Triton X-100, 2 mM EDTA, 20 mM Tris-HCl), and twice with TE buffer (1 mM EDTA, 10 mM Tris-HCl). DNA recovery and purifications were performed as for Histone ChIP. Results in Fig. 6B were normalized to IGN5.

### Computational model

For a detailed description of the computational model, its assumptions, implementation, and analysis of simulation output, see Supplementary Information. Additional experimental considerations for modelling are also presented in the Supplementary Information.

## Acknowledgements

The authors would like to thank members of the Dean and Howard labs for thoughtful inputs and Shuqin Chen for excellent technical assistance. This work was funded by European Research Council Advanced Grant (EPISWITCH, 833254), Wellcome Trust (210654/Z/18/Z), and a Royal Society Professorship (RP\R1\180002) to CD, and BBSRC Institute Strategic Programmes (BB/J004588/1 and BB/P013511/1) and EPSRC/BBSRC Physics of Life grant (EP/T00214X/1) to MH and CD. Eduardo Mateo-Bonmatí would like to thank grant RYC2021-030895-I, funded by MCIN/AEI and by European Union NextGenerationEU/PRTR. Svenja Reeck was funded by the HORIZON EUROPE Marie Sklodowska-Curie Actions PEP-NET (Predictive Epigenetics: Fusing Theory and Experiment) grant 813282.

## Figure Captions

**Figure 4 – Supplementary 4A:**
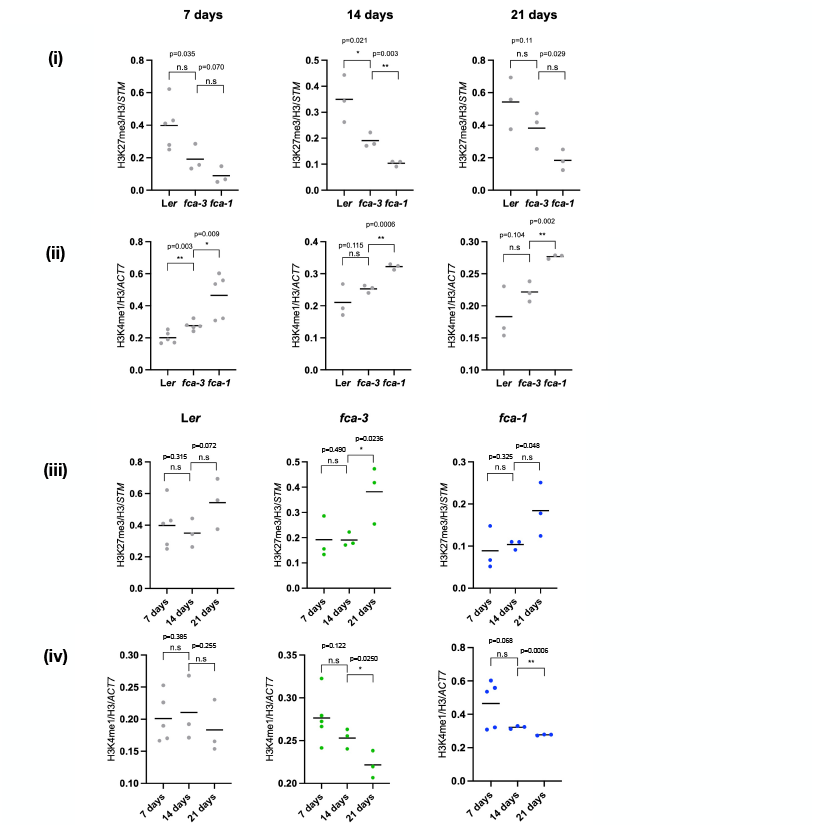
Statistical comparisons of average ChIP-qPCR measured H3K27me3 (i,iii) and H3K4me1 (ii,iv) levels across the locus, between genotypes (i, ii) and between timepoints (iii,iv). In all cases the data points show the measured modification level averaged across 15 primers covering the *FLC* locus (full modification profiles shown in Fig. 4A,B). Horizontal bars indicate mean of replicates. Statistical comparisons for the experimental data were carried out using the Student’s t-test and corrected for multiple comparisons using the Holm-Sidak approach (*, **, *** indicate significance thresholds of 0.05, 0.01, and 0.001 respectively).

**Figure 4 – Supplementary 4B:**
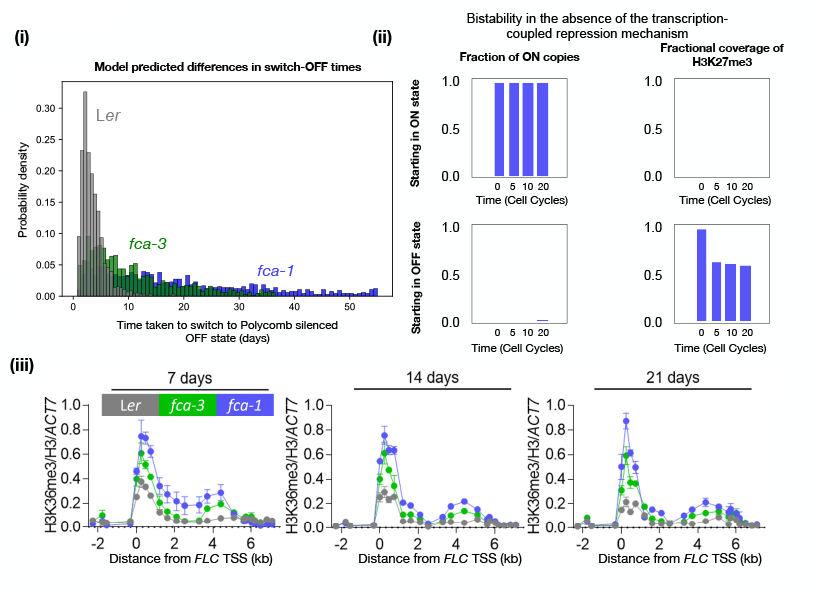
**(i)** Histograms showing probability distribution of switch OFF times for an individual locus, for L*er, fca-3*, and *fca-1*. Histograms are generated by simulating 1000 copies, started in a full H3K4me1 covered ON state, and simulated over 60 cell cycles. **(ii)** Model predicts that the locus is in a bistable regime in the absence of the proximal termination mechanism (proximal termination probability set to residual value – see model description in Supplementary Information and parameter definitions/values in Supplementary Table 3), with both ON state and OFF state stably maintained across several cell cycles. Plots show fraction of ON copies (left column) and H3K27me3 fractional coverage (right column) for 100 simulated loci started in an ON state (full H3K4me1 coverage) (top row) or started in an OFF state (full H3K27me3 coverage) (bottom row). Fractions are shown at the starting timepoint (0), and at the end of 5, 10, and 20 cell cycles from the start. During steady state maintenance of the OFF state, the cell cycle-averaged H3K27me3 stabilises at an intermediate level due to the periodic dilution by DNA replication and subsequent slow accumulation. **(iii)** ChIP-qPCR time course measurements of H3K36me3 across the *FLC* locus in the parental L*er* genotype and the *fca* mutants. Each dot represents an amplicon. Error bars represent mean +/- s.e.m. (n = 3 biological replicates). H3K36me3 levels are shown normalised to H3 and to the control locus *ACT7*.

**Figure 4 – Supplementary 4C:**
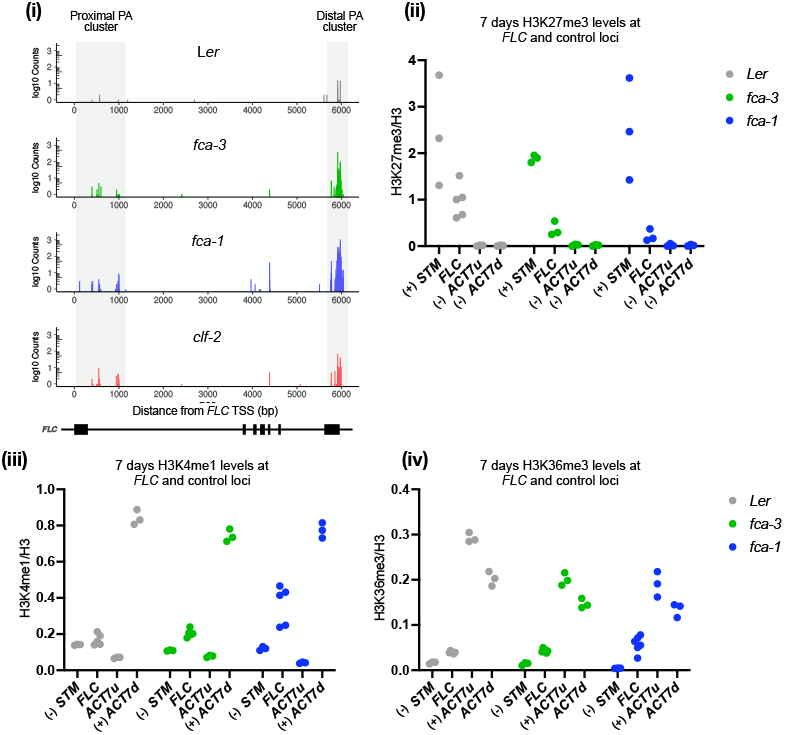
**(i)** Quantseq data for 7-day old seedlings (7 days after sowing), showing quantification of polyadenylated transcripts mapped to the *FLC* locus in the parental Landsberg (L*er*) genotype, the *fca* mutants (L*er* and *fca* mutants data same as shown in Fig. 2B), and the PRC2 methyltransferase mutant *clf-2*. **(ii)** ChIP-qPCR measured H3K27me3 levels (normalised to H3) shown for the three *fca* genotypes at *FLC* as well as at the designated positive (+) and negative control loci (-) for this modification. **(iii)** ChIP-qPCR measured H3K4me1 levels (normalised to H3) shown for the three *fca* genotypes at *FLC* as well as at the designated positive (+) and negative control loci (-) for this modification. **(iv)** ChIP-qPCR measured H3K36me3 levels (normalised to H3) shown for the three *fca* genotypes at *FLC* as well as at the designated positive (+) and negative control loci (-) for this modification.

**Figure 4 – Supplementary 4D:**
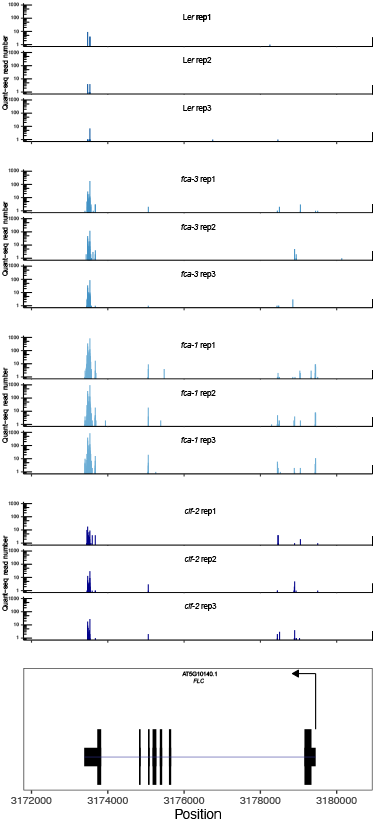
Individual replicates for Quantseq data shown in Fig. S4C.

**Figure 5 – Supplementary 5A:**
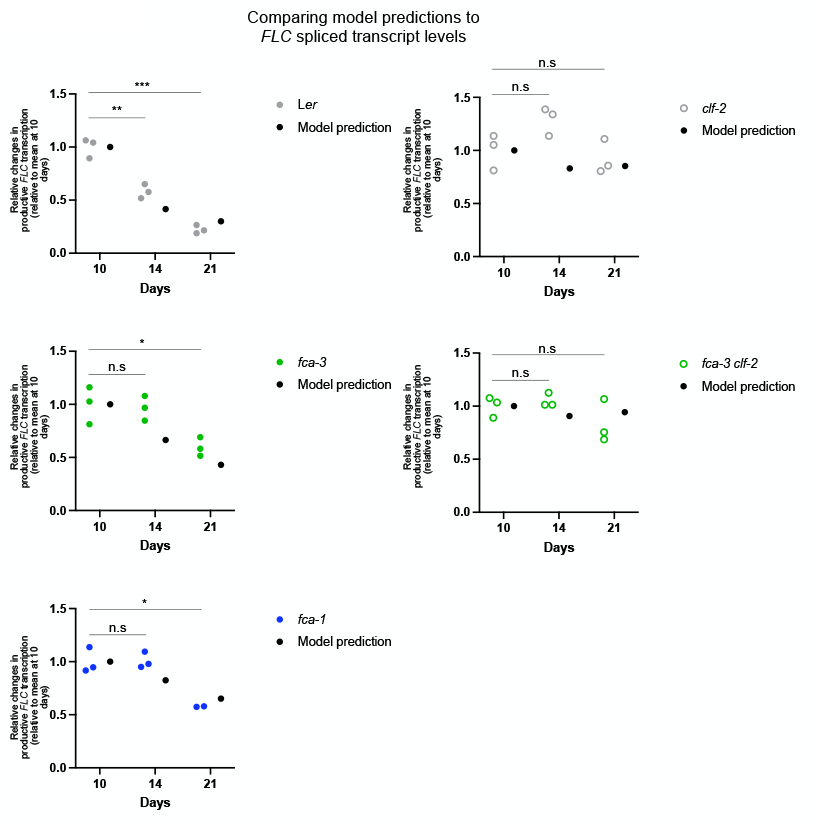
Changes in Spliced *FLC* transcript level. Comparison of model predicted changes in *FLC* transcription, with whole seedling *FLC* spliced transcript levels measured by qPCR in the *fca* genotypes (L*er, fca-3, fca-1*) and the *clf* mutants (*clf-2* and *fca-3 clf-2* double mutant). Experimental data is shown normalised to measured levels for *UBQ10*. Statistical comparisons for the experimental data were carried out using the Student’s t-test and corrected for multiple comparisons using the Holm-Sidak approach (*, **, *** indicate significance thresholds of 0.05, 0.01, and 0.001 respectively). Model predictions correspond to the predicted frequency of *FLC* sense distal transcription events at the corresponding timepoints, averaged over 1000 simulated loci. Simulations for L*er, fca-3, fca-1* are identical to those used to predict changes in histone modification levels (Fig. 4). Simulations for the *clf* mutants (*clf-2* and *fca-3 clf-2*) use changed parameter values for reactions involving H3K27me3 addition by PRC2 – see Supplementary Information and Supplementary Table 2. For comparison of reltive changes, both model predictions and data are normalised to the level at 10 days.

**Figure 5 – Supplementary 5B:**
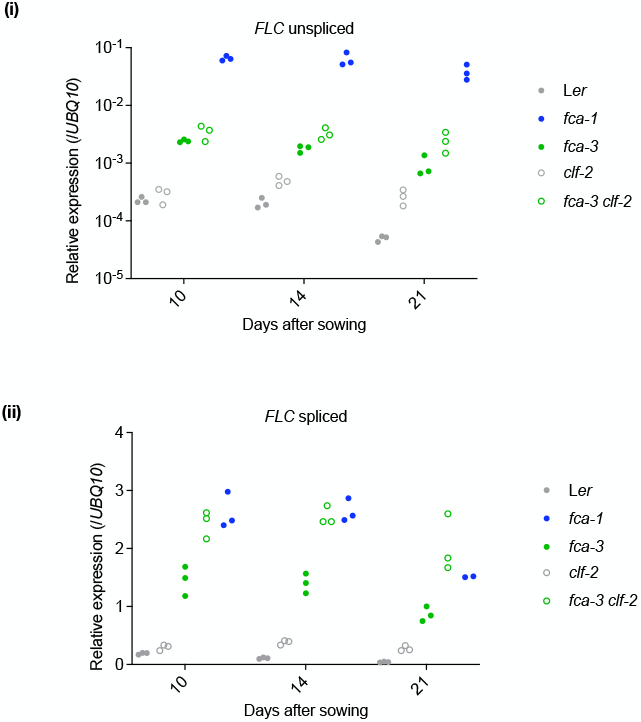
**(i)** *FLC* unspliced transcript levels in whole seedlings measured by qPCR. **(ii)** *FLC* spliced transcript levels in whole seedlings measured by qPCR. Data is shown normalised to the measured levels for *UBQ10*.

**Figure 5 - Supplementary 5C:**
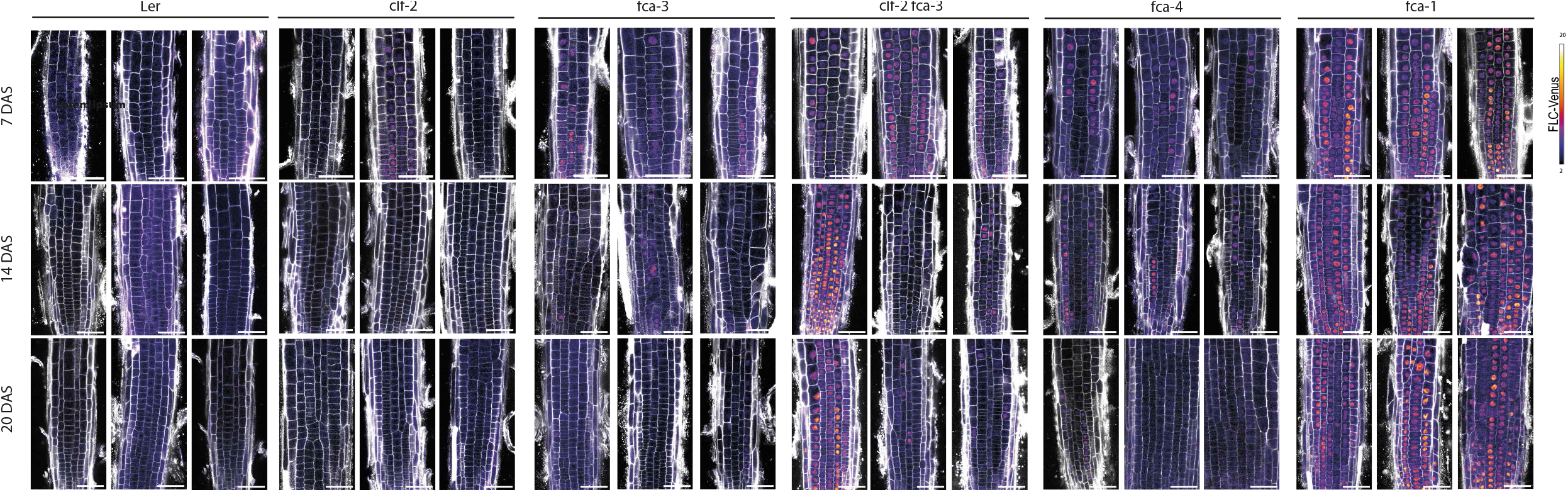
FLC-Venus time course replicates with additional genotypes. Additional representative images of FLC-Venus signal in Ler, *clf-2, fca-3, fca-3 clf-2* as well as an additional intermediate allele *fca-4* and the strong allele *fca-1* at three time points. FLC-Venus intensity is indicated by colour map; grey shows the cell wall. Scale bar, 50 □ μm. DAS stands for days after sowing.

**Figure S6:**
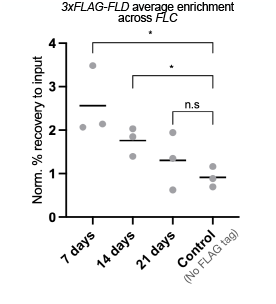
Statistical analysis of changes over time in 3xFLAG-FLD enrichment across the locus. Data points indicate average measured level over 16 primers covering the *FLC* locus (full dataset shown in Fig. 6B). Horizontal bar indicates mean over replicates. Statistical comparisons for the experimental data were carried out using the Student’s t-test and corrected for multiple comparisons using the Holm-Sidak approach (*, indicates a significance threshold of 0.05).

## Graphical Abstract

Schematic showing the general interplay between transcriptional activity, chromatin state, and co-transcriptional processing that controls the transcriptional state, and the specific aspects of this interplay that regulate transcription at the *FLC* locus.

## Supplementary Information

### Computational Model

We built a stochastic model describing the dynamics of the activating and repressive histone modifications of interest, across a single *FLC* locus. The model also describes transcription events at the locus – sense and antisense – as well as alternative polyadenylation during these transcription events.

### Model features

#### Activating and repressive histone modifications

The mathematical model describes the modification state of each H3 histone at the *FLC* locus, with the whole locus represented by an array of 30 nucleosomes, consistent with the approximately 6 kilobases extent of the locus (nucleosomal DNA + linker DNA spans approximately 200 bp). The modifications described are: (1) H3K4me1, an activating modification associated with gene bodies in Arabidopsis, and known to exhibit significant differences between active and silenced states of *FLC*; (2) the Polycomb-mediated repressive modification H3K27me3 as well as possible H3K27me2, H3K27me1 and me0 states. The model assumes mutual exclusivity of K4me and K27me on the same H3 tail in line with experimentally observed mutual exclusivity in mESCs (Shema et al., 2016; Voigt et al., 2012) (Kim et al., 2013; Schmitges et al., 2011). Evidence from mammalian and yeast studies indicate that H3K27me and H3K4me are added non-processively (McCabe et al., 2012; Soares et al., 2017). Therefore, the model assumes stepwise addition of these modifications.

#### Sense and Antisense transcription

The model describes sense and antisense transcription events at the locus. The frequency of these events (assumed to represent transcription initiation or transition to elongation) is determined by the coverage of repressive H3K27me3 across the locus. This is consistent with Polycomb (PRC1) binding to H3K27me3 and controlling accessibility to gene regulatory elements by mediating compaction (Blackledge and Klose, 2021), as well as other potential H3K27me3-mediated mechanisms, such as inhibition of RNA Pol II release from the Transcription Start Site (TSS) (Zhang et al., 2020), thus preventing transitions into an actively transcribing Pol II state (Brookes et al., 2012). Our data indicates that both sense and antisense transcription at *FLC* are silenced by Polycomb – both types of transcripts are present at higher levels in the absence of Polycomb silencing, as observed in genotypes containing the *FLC* activator FRIGIDA, and in loss of function mutants of FCA and FLD.

#### Co-transcriptional addition of activating modifications

Activating modification such as H3K4me and H3K36me in many contexts and across organisms are known to be added co-transcriptionally (Chan et al., 2022). The association of the H3K4 methyltransferase complex with elongating RNA Pol II and the addition of this modification during transcriptional elongation has been demonstrated in *S. cerevisiae* (Soares et al., 2017). In the case of Arabidopsis H3K4me1, a genome wide analysis of the localization patterns of H3K4me1, of RNA Pol II, and of the different H3K4 methyltransferases, indicates that the co-transcriptional addition of H3K4me1 over the gene body is mediated mainly by the methyltransferase ATXR7 (Oya et al., 2022). Consistent with this role, an *ATXR7* mutant has been shown to partially suppress *FLC* activation by the activator FRIGIDA (Tamada et al., 2009).

#### Alternative polyadenylation mediated by coverage of activating modifications

The model assumes that higher overall coverage of H3K4me1 across the locus promotes distal termination of transcription. This assumption is in line with a potential feedback mechanism where H3K4me1 enables binding of the H3K36me3 methyltransferase (SDG8) and consequent H3K36me3-mediated assembly of splicing factors (Leung et al., 2019) which facilitate co-transcriptional processing. Such a role for H3K4 methylation is also consistent with evidence in yeast (Soares et al., 2017) and mammalian cells (Wang et al., 2023), indicating a role in facilitating productive transcriptional elongation.

#### Proximal termination mediated removal of activating modifications

The model assumes proximal termination of both sense and antisense mediates removal of H3K4me1. This is in line with the potential function of FLD as a demethylase (Inagaki et al., 2021), and the fact that genetically, FLD relies on FCA-mediated proximal termination of antisense (Liu et al., 2007). The removal of H3K4me1 resulting from proximal PA is assumed to take place uniformly across the whole locus, as the data we currently have does not allow us to observe the spatial details of this process.

#### Polycomb nucleation and spreading

The model describes nucleation of H3K27me by Polycomb in the *FLC* nucleation region, where Polycomb silencing is known to be nucleated during cold-induced chromatin silencing of *FLC*. Subsequent H3K27me3 spreading across the locus in this context is known to depend on the plant PRC2 methyltransferase CLF and on an active cell cycle, while nucleation does not depend on either. Experimental evidence from a genome-wide ChIP-seq study showing H3K27me3 patterns in a *clf* mutant (Shu et al., 2019) indicates that a similar mechanism is involved in the early developmental silencing of *FLC* – the *clf* mutant shows a clear H3K27me3 peak in the *FLC* nucleation region, while spreading is compromised. Consistent with this, in the wild-type, H3K27me3 is assumed to spread to the rest of the locus through interactions with the nucleation region, through read-write feedback interactions. The spreading is assumed to rely on an active cell cycle – in the model, this is achieved by having the interactions with the nucleation region stronger in S/G2 phases.

#### Looping interactions mediating Polycomb activity

Experimental evidence indicates the existence of a gene loop mediating an interaction between the 5’ and 3’ ends of *FLC* (Crevillen et al., 2013; Li et al., 2018), with the level of interaction potentially depending the transcriptional state of the locus (Li et al., 2018). Significantly, this looping interaction is present in the silenced *FLC* state in the reference genotype Col-0, and consistently, a secondary peak of H3K27me3 is observed in published ChIP-seq data in *clf* (Shu et al., 2019). Therefore, we capture this interaction in the model by allowing for stronger interactions with the nucleation region for a designated “looping region” within the gene body.

#### Productive transcription antagonises H3K27me

Experimental evidence from mammalian systems indicates that productive transcription can promote removal of H3K27me through histone exchange. In addition, the H3K27me demethylases have been shown to physically interact with and co-localise on chromatin with elongating RNA Pol II, in *Drosophila* (Smith et al., 2008) and more specifically at the *FLC* locus (Yang et al., 2016). Both features have been captured in previous models of Polycomb silencing (Berry et al., 2017), and they are also incorporated in the current model. These features allow the rate of H3K27me turnover across the locus to be determined by the frequency of transcription events. Our model also incorporates alternative PA of both sense and antisense transcription, allowing proximal PA and termination of RNA Pol II. The model assumes that only distal PA and termination (i.e., productive transcription events) can remove H3K27me through the above mechanisms.

#### Mutual exclusivity of sense and antisense at the same locus

We previously found that distal antisense transcription and sense *FLC* transcription are mutually exclusive at individual *FLC* loci (Rosa et al., 2016). Consistent with this mutual exclusivity, we have observed an anticorrelation between the levels of distal antisense transcripts and sense transcripts in multiple contexts, including different chromatin states of the *FLC* locus (Ietswaart et al., 2017; Zhao et al., 2021). While this evidence demonstrates a mutual exclusivity between full length (distal) sense and antisense transcription events, we currently do not have data to determine whether the mutual exclusivity applies to proximally terminated transcription events. However, for simplicity, the model assumes mutual exclusivity for both types of transcription events. This is realised by having a refractory period after a sense (antisense) transcription event, during which antisense (sense) initiation is not allowed to occur. The model assumes equal refractory periods for sense and antisense transcription, as well as for distal and proximal termination.

#### Inheritance of modified histones during DNA replication

In line with recent evidence indicating that only H3-H4 tetramers carrying repressive modifications are reliably inherited across DNA replication (Escobar et al., 2021), the model assumes that H3 pairs corresponding to an individual nucleosome are inherited with probability half to a daughter strand, only if at least one of the H3 carries K27 methylation.

#### Additional considerations for the model

##### Absence of a role for the L*er Mutator*-like transposable element at *FLC*

The *FLC* allele in the L*er* accession is known to contain a *Mutator*-like transposable element (TE) in intron 1 that causes lower expression from this allele relative to *FLC* alleles in other accessions (Michaels et al., 2003). This lower expression results from RNAi-mediated chromatin silencing of the TE element (Liu et al., 2004). However, experimental evidence indicates that the intermediate *FLC* expression levels in *fca* mutants are unlikely to be mediated by the TE. The TE has been shown to act *in cis* (Michaels et al., 2003), as expression of an *FLC* allele from the Columbia-0 (Col-0) accession is unaffected in F1 plants generated by a L*er* to Col-0 cross, while the L*er FLC* allele expression is attenuated by the TE. In our case, the intermediate expression in *fca* mutants could be observed for both the endogenous *FLC* (L*er FLC* containing TE) and the transgenic Venus tagged *FLC* (Col-0 *FLC* sequence), arguing against a role for the *cis*-acting TE mechanism in generating intermediate transcription states.

#### Role for H3K36me3

The above model only considers the histone modifications H3K4me1 and H3K27me, and does not explicitly consider H3K36me3. As discussed in the text, the Arabidopsis H3K36 methyltransferase SDG8 is known to recognise H3K4me1, thus mechanistically linking the two modifications. Combined with the H3K36me3 requirement for effective co-transcriptional splicing and recruitment of splicing factors as reported in *S. cerevisiae* (Leung et al., 2019) this suggests a mechanism where FLD-mediated removal of H3K4me1 may cause reduced H3K36 methylation, which in turn promotes further early transcription termination by disrupting co-transcriptional splicing. For simplicity, the model omits the H3K36me3 step and assumes that H3K4me1 implicitly mediates this effect on termination.

#### Direct physical interaction between transcription-coupled repression and Polycomb

FCA has been reported to interact directly with the Arabidopsis PRC2 methyltransferase CLF *in vivo* (Tian et al., 2019). However, we have found no evidence of this interaction in our FCA immunoprecipitation followed by mass spectrometry (IP-MS) data or other published CLF IP-MS data (Fang et al., 2019; Liang et al., 2015), and hence this interaction is not further considered in our model or other analyses.

### Model implementation

#### Locus description

The whole locus represented by an array of 30 nucleosomes (60 H3 histones), consistent with the approximately 6 kilobase extent of the locus (nucleosomal DNA + linker DNA spans approximately 200 bp).

#### Mutually exclusive H3 modifications

Each H3 histone can be in one of five modification mutually exclusive states: {-1,0,1,2,3} representing H3K4me1, H3 unmodified on K4 and K27, H3K27me1, me2, and me3 respectively. This means addition of K27me can take place only in the absence of H3K4me1 and vice versa. All transitions are assumed to be non-processive: only transitions involving addition/removal of a single methyl group are allowed. The modification state of the locus is representing by an array S representing the modification states of the 60 histones. Each entry *S*_*i*_ can take the values {-1,0,1,2,3}. The array S is used to store the current state of all 60 histones over the course of the simulation. Note that we do not describe H3K36me3 in the model, as the evidence indicates it is downstream of H3K4me1, as described above and in the main text.

#### Alternative termination of transcription

Each transcription event (sense or antisense) has a probability of being proximally terminated. This probability is determined by both the current H3K4me1 fractional coverage (whole locus) and the FCA effectiveness parameter, as follows:

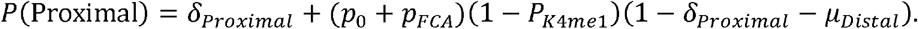

The probability of distal termination is given by:

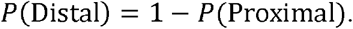

Here *p*_*FCA*_ represents FCA effectiveness parameter and *p*_0_ captures possible FCA independent proximal termination. (*p*_0_ + *p*_*FCA*_)is constrained to be less than or equal to 1. *P*_*K*4*me*1_ represents the fractional coverage of H3K4me1, *δ*_*Proximal*_ is a residual proximal termination probability (some proximal termination occurs even when *P*_*K*4*me*1_=1), and *μ*_*Distal*_ is a residual probability for distal termination (some distal termination occurs even when *p*_*FCA*_=1 or *P*_*K*4*me*1_=0).

#### Transcriptional antagonism of Polycomb

(1) Histone exchange: during each distally terminated transcription event, iterating over the histones (entries of array S), for each histone, we generate a pseudo-random number from a uniform distribution (0,1) and set both entries corresponding to the current nucleosome to 0 if the generated number is less than *p*_*ex*_/2 (see Supplementary Table). (2) Co-transcriptional Demethylation: During each distally terminated transcription event, iterating over the entries of array *S*, for each histone, we generate a pseudo-random number from a uniform distribution (0,1) and reduce the methylation state by one if the current state is {1, 2, or 3} AND the generated number is less than *P*_*dem*_ (see Supplementary Table).

#### Transcription mediated effects on H3K4 methylation

(1) Co-transcriptional addition of H3K4me1: Each unmodified H3 histone has a fixed probability 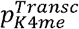, of being methylated during a productive (distally terminated) transcription event (not in case of proximal termination). (2) H3K4 demethylation by FLD: Each H3 histone carrying H3K4me1 has a fixed probability 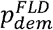 of losing this modification during a proximally terminated transcription event.

#### H3K4me1 dynamics

The background turnover rate (independent of proximal termination and FLD) of H3K4me1 is set so that overall turnover rates of H3K27me3 and H3K4me1 are in the ratio 3.128/0.959, as measured in HeLa cells by a SILAC technique in (Zee et al., 2010). The H3K4 methylation propensities (background and transcription mediated) are chosen to be significantly higher than the K27me3 propensities, consistent with faster H3K4 methylation seen in time course evidence from mammalian systems.

#### H3K27me3 dynamics

Values for methylation rate constants are mostly chosen as in (Berry et al., 2017). The overall rate β alone is adjusted from its value in (Berry et al., 2017) to allow for a changed interaction paradigm – instead of just nearest neighbour interactions, the current model allows long ranged interactions between nucleosomes across the whole locus, with any nucleosome being able to interact with any other within the locus at some rate (see below). To account for nucleation, spreading and the looping interaction, the nucleosome array representing the whole locus is divided into three subsets: nucleation region (NR, nucleosomes 1 to 3, i.e., histones 1 to 6), looping region (LR, nucleosomes 20 to 22, i.e., histones 39 to 44), and body region (Body, remaining nucleosomes). The choice of nucleosomes 19 to 21 for the looping region is consistent with the observed secondary nucleation peak observed around 4kb downstream of the *FLC* TSS in published data for the *clf* mutant (Shu et al., 2019).

The methylation propensities at each histone *i* are computed as:

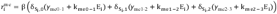

where *E*_*i*_ represents the PRC2 read-write feedback contribution.

For nucleation region histones, the PRC2 read-write feedback mediated contribution to the H3K27 methylation propensity is computed as:

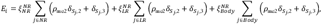

where *ρ*_*me2*_ is the relative activation of PRC2 by H3K27me2 as compared to H3K27me3. For the looping region histones, the PRC2 read-write feedback mediated contribution to the H3K27 methylation propensity is computed as:

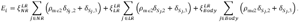

For the remaining gene body histones, the PRC2 read-write feedback mediated contribution to the H3K27 methylation propensity is computed as:

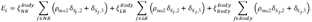

Here the parameters *ξ* are chosen so that interactions between nucleosomes within the nucleation and looping region histones produce the highest contribution to feedback methylation, followed by the contribution of NR and LR nucleosomes to methylation of other gene body nucleosomes. This is consistent with the PRC2 enzymatic machinery localising mainly to the nucleation region. Also consistent with this, we assume lower contributions from gene body histones to the methylation of NR and LR nucleosomes, and the lowest contribution from interactions between gene body histones.

Hence, we set the above parameters as follows:

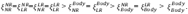

Following (Berry et al., 2017), a background “noisy” removal of H3K27 methylation is also included in the model, represented by the rate constant *γ*_*dem*_.

#### Dependence on an active cell cycle

To implement the dependence of H3K27me3 spreading on an active cell cycle (as observed experimentally in (Yang et al., 2017)), the total cell cycle duration is divided into two parts: G1 and S/G2/M, with replication of the locus occurring at the midpoint of S/G2/M. The duration of G1 is assumed to be approximately half the total duration of the cell-cycle. The four parameters corresponding to interactions between the NR/LR and the body region 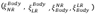 are reduced by a factor of 10 during the part of the cell cycle corresponding to G1 phase.

#### H3K27me3 spreading mutant

For simulating the H3K27me3 spreading mutant (*clf-2*), we lower the read-write feedback contributions from the NR and LR nucleosomes to methylation of gene body nucleosomes (i.e., lower 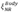 and 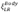), as well as the contribution from the gene body nucleosomes to methylation of the NR and LR nucleosomes (i.e., lower 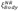 and 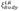). Since the loss of one of the PRC2 methyltransferases may also be expected to partially affect H3K27me3 addition within the nucleation region itself, we also slightly reduce the feedback contributions between nucleosomes inside the NR and LR regions (i.e., lower 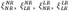). These parameter values are specified in supplementary Table 2.

#### De novo addition of H3K27me3

In addition to the differences in PRC2 feedback mediated contributions, the nucleation region (first three nucleosomes) is assumed to have a significantly higher *de novo* contribution to the H3K27 methylation propensity (*γ*_*me*_ parameters), consistent with evidence for *de novo* PRC2 targeting to this region during early developmental silencing of *FLC* (Shu et al., 2019).

#### H3K27me3 fractional coverage measure

Our previous studies of *FLC* silencing indicate that nucleation region H3K27me3 alone can significantly repress transcription, with the body region H3K27me3 making a relatively smaller additional contribution to silencing. This finding is different to the assumptions of Berry et al., where both H3K27me2 and H3K27me3 could repress. Furthermore, nucleation of H3K27me3 in the cold is sufficient to switch the locus to a digitally OFF state, as seen by FLC Venus imaging in the roots (Lovkvist et al., 2021). Potentially different contributions of the nucleation and looping region H3K27me3 versus the body region H3K27me3 to the transcriptional silencing are captured by allowing a higher contribution from these NR and LR nucleosomes when computing the overall fractional coverage of H3K27me3:

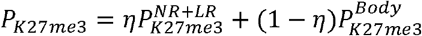

Here 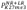 and 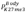 represent fractional coverage of H3K27me3 in the nucleation region/looping region and the rest of the locus, respectively, and 0< *η* < 1. The overall fractional coverage, *P*_*K*27*me*3_ computed above, is then used to determine the transcription propensity (see below).

#### Polycomb silencing of transcription

H3K27me3 coverage across the locus is assumed to determine the “transcription initiation” frequency. This frequency is assumed to be a piecewise linear function of H3K27me3 coverage with a saturation threshold (similar to the model in (Berry et al., 2017)). Note that this frequency can also represent the transition of initiated Pol II into the elongating phase. Nucleation region and looping region contributions to the fractional coverage are assumed to be higher than the rest of the locus (see above). H3K27me3 coverage also has the same effect on sense and antisense initiation frequency.

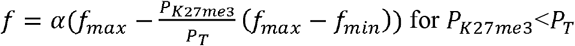

and

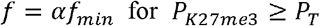

We also allow for a slightly higher transcription frequency in the H3K27me3 nucleated state compared to a fully H3K27me3 spread state. This is consistent with our previous datasets, where *FLC* transcriptional output is observed to reduce further during the post-cold transition to a spread state following nucleation. To enable this behaviour, we set *η* < *P*_*T*_ so that the full silencing threshold is not reached even with full coverage of H3K27me3 in the nucleation and looping regions (i.e., even with 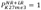).

Published evidence based on sequencing of nascent RNA as well as our own qPCR data (Wu et al., 2016) indicates lower frequency of antisense transcription compared to sense transcription. This difference between sense and antisense transcription is captured by using different values of α for sense and antisense initiation frequency. Note that H3K4me1 does not affect the initiation frequency.

#### Histone inheritance during DNA replication

For each individual simulation, the DNA replication times are fixed deterministically, assuming a cell cycle duration of 22h (Rahni and Birnbaum, 2019). At each DNA replication timepoint, the Gillespie Stochastic Simulation Algorithm (SSA) is interrupted. We then iterate over the entries of the array S, where for each nucleosome at the locus carrying at least one H3K27me, its modification state is set to 0, i.e., the pair of entries corresponding to the nucleosome is set to 0, with a probability of 0.5. We assume here random inheritance of intact H3-H4 nucleosomes, with equal probability to each daughter strand. For each nucleosome, this is done by generating a pseudo-random number from a uniform distribution (0,1) and setting the corresponding histone modification states to 0 if the generated number is less than 0.5. In contrast, for each nucleosome carrying no H3K27me, the modification state is always set to zero, i.e., no inheritance of nucleosomes carrying only H3K4me1.

### Simulations

Model simulations were carried out using the Gillespie exact SSA (Gillespie, 1977). Simulations were coded in C++ and compiled using Clang/LLVM (version 14.0.0). The Mersenne Twister 19937 algorithm (Matsumoto and Nishimura, 1998), implemented in the GNU Scientific Library (GSL version 2.7), was used to generate pseudorandom numbers for the Gillespie SSA. The simulation code can be accessed at: https://github.com/gm1613/FLC_transcription_coupled_repression_model.git

### Initial conditions

In all cases – both for the analog module in isolation and the full model, the initial state was set to full coverage of H3K4me1.

### Analysis of simulation output

Average levels of fractional coverage of histone modifications and average frequency of different transcription events and termination site usage (sense distal, antisense distal, sense proximal, antisense proximal) were computed by averaging over 500 individual trajectories. Individual trajectories were started at random (uniformly distributed) times in the cell cycle to avoid having all trajectories synchronised. To quantify switching to the digital OFF state (which in the model can correspond either to a Polycomb nucleated state or a spread state), we have to define what qualifies as an OFF state. In our analysis we classify an OFF state as any state for which:

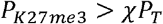

where the parameter *χ* is chosen so that *χP*_*T*_ < *η*, and all the other parameters/quantities are as defined above. This condition on *χ* ensures that an H3K27me3 build-up in the nucleation region (even if this modification is absent across the rest of the locus), can be sufficient to qualify as a digitally OFF state. In all simulation results shown, *χ* is set to 0.25.

### Comparing simulation output to experimental data

#### Comparison to timecourse ChIP-qPCR data

In Fig. 4 we compare the model predicted levels of H3K27me3 and H3K4me1 to the normalised levels measured by ChIP-qPCR (averaged over primers across the locus). The model predicted levels consist of an average over multiple simulated copies (1000 copies in all cases). While the model only describes a set of *FLC* copies in dividing cells, the ChIP-qPCR measurement is performed on whole-seedling tissue. Therefore, we compare a model predicted population average to the experimental data. Note that we expect there to be heterogeneity in the modification levels between tissues, and differences in the dynamics between dividing and non-dividing cells. We have previously addressed these differences at *FLC* in detail using a population level Ordinary Differential Equation model of *FLC* copies switching between different states (Questa et al., 2020). However, capturing these aspects is beyond the scope of the current study. We therefore make a simpler assumption: that at all timepoints and across genotypes, there exist two subpopulations of *FLC* copies (at the whole plant level) in addition to the population of copies represented in the model. We assume that one of these subpopulations consists entirely of active *FLC* copies (with high H3K4me1 coverage) with no possibility of PRC2 silencing, and the other subpopulation consists of PRC2 silenced copies (with high H3K27me3 coverage), with no possibility of switching to an active state. We assume that these copies form a fixed fraction of the total *FLC* copies, fixed across timepoints and across genotypes (hence independent of FCA). Note that these subpopulations must also consist of dividing cells in order for their fractions to be maintained constant over time. We do not explicitly simulate these subpopulations. Instead, we account for their contribution to the averaged histone modification levels. The contribution from each copy in the active subpopulation to H3K4me1 is assumed to be the same as the average steady state level at non-Polycomb silenced copies in the *fca-1* (null mutant) model, and with no contribution to the H3K27me3 level. Similarly, the contribution from each copy in the silenced subpopulation to H3K27me3 is assumed to be the same as the average level at PRC2 silenced copies predicted by the L*er* model (these levels are similar in the L*er, fca-3, and fca-1* models). We assume that these silenced copies make no contribution to the H3K4me1 level. The sizes of these two subpopulations relative to the population of copies represented by the model form two model parameters that we change to fit the data. The fits presented in Fig. 4C,D assume that the active subpopulation is 0.2 times the size of the modelled population, while the silenced subpopulation is 0.1 times the size of the modelled population.

#### Comparison to ChIP profile across *FLC*

In Fig. 5E, we compare the model predicted H3K27me3 profiles at *FLC* in the L*er* parental genotype and the Polycomb spreading mutant *clf-2* to the ChIP-qPCR measured profiles in these genotypes. Note that the model predicted profile is an average across multiple simulated copies (1000 copies). To do this we used an approach developed in (Wu et al., 2016) for comparing model predicted Pol II profiles to those measured by ChIP. In our case, this method generates a predicted H3K27me3 ChIP profile by combining the fully spatially resolved model predicted profile with the resolution limiting effects arising from the sonication generated fragment size distribution inherent in ChIP based measurements. Mathematically, this method involves a convolution of the full model predicted profile with the probability distribution of fragment sizes generated during the ChIP experiment. Here we use the same fragment size distribution estimated in (Wu et al., 2016). The code used to carry out this analysis is also provided at: https://github.com/gm1613/FLC_transcription_coupled_repression_model.git

##### Comparison of predicted *FLC* shutdown

<u>In Fig. 5C and Fig. S5A, we compare the model predicted</u> *<u>FLC</u>* <u>transcriptional shutdown over time to changes in</u> *<u>FLC</u>* <u>unspliced (Fig. 5C)and</u> *<u>FLC</u>* <u>spliced (Fig. S5A) RNA levels, measured by qPCR at the whole plant level. The model predicted</u> *<u>FLC</u>* <u>transcriptional output is computed as the average frequency of</u> *<u>FLC</u>* <u>sense distal transcription events (transcription events per hour). This frequency is computed by averaging sense distal transcription events over a 6hr time window (an approximation based on our estimate of ∼6hr</u> *<u>FLC</u>* <u>mRNA half-life) leading to the respective sampling timepoints (to match the experiments, we choose 10 day, 14 day, and 21 day timepoints). We note that the model is not designed to capture absolute RNA levels in the different genotypes, since this may depend on differences in their stability, as well as heterogeneity between tissue types which are not described in the model. What the model is intended to capture, is the quantitative trends, including differences between genotypes and changes over time, i.e. a reduction over time in some genotypes which may be disrupted in others. A whole-plant level measurement of</u> *<u>FLC</u>* <u>RNA is an appropriate quantity for capturing these types of differences, and we therefore use these measurements for comparison to the model predictions. Since the model does not capture absolute RNA levels, for comparison of experimental timecourse data to model predictions, we consider changes (i.e., fold changes) relative to the first timepoint (10 days).</u>

**Supplemental Table S1.**
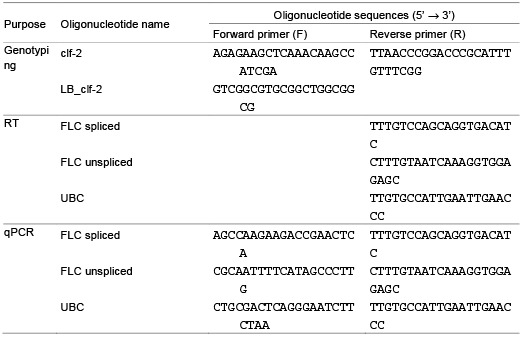
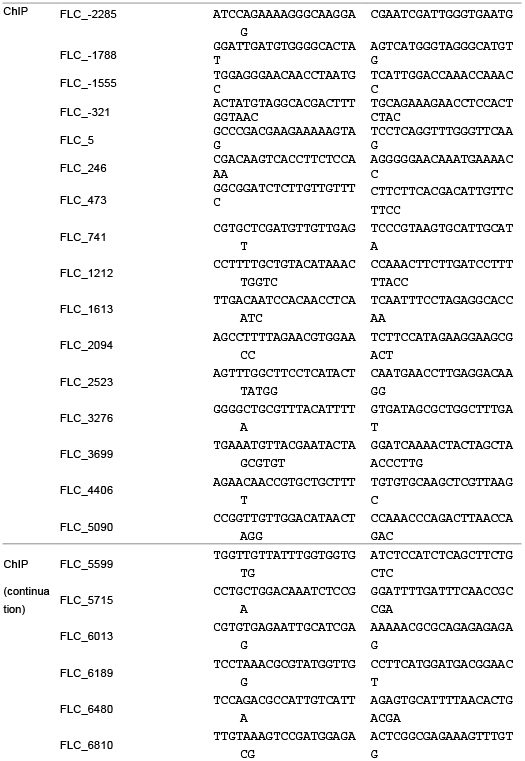
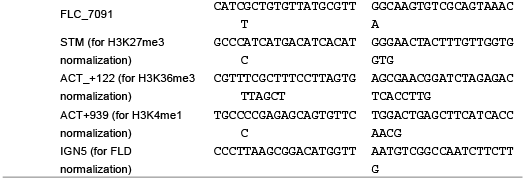
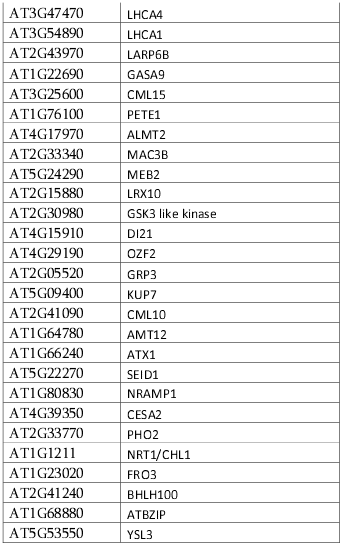
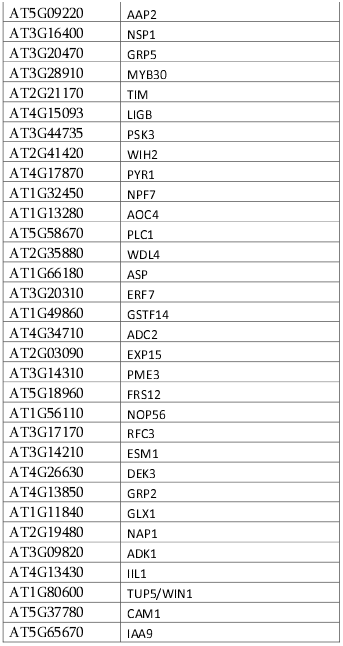
Primer sets used in this work.

